# Elastohydrodynamical instabilities of active filaments, arrays and carpets analyzed using slender body theory

**DOI:** 10.1101/2020.03.10.986596

**Authors:** Ashok S. Sangani, Arvind Gopinath

**Affiliations:** Department of Biomedical and Chemical Engineering, Syracuse University, Syracuse, NY 13244 USA; Department of Bioengineering, University of California, Merced, CA 95343 USA

## Abstract

The rhythmic motions and wave-like planar oscillations in filamentous soft structures are ubiquitous in biology. Inspired by these, recent work has focused on the creation of synthetic colloid-based active mimics that can be used to move, transport cargo, and generate fluid flows. Underlying the functionality of these mimics is the coupling between elasticity, geometry, dissipation due to the fluid, and active force or moment generated by the system. Here, we use slender body theory to analyze the linear stability of a subset of these - active elastic filaments, filament arrays and filament carpets - animated by follower forces. Follower forces can be external or internal forces that always act along the filament contour. The application of slender body theory enables the accurate inclusion of hydrodynamic effects, screening due to boundaries, and interactions between filaments. We first study the stability of fixed and freely suspended sphere-filament assemblies, calculate neutral stability curves separating stable oscillatory states from stable straight states, and quantify the frequency of emergent oscillations. When shadowing effects due to the physical presence of the spherical boundary are taken into account, the results from the slender body theory differ from that obtained using local resistivity theory. Next, we examine the onset of instabilities in a small cluster of filaments attached to a wall and examine how the critical force for onset of instability and the frequency of sustained oscillations depend on the number of filaments and the spacing between the filaments. Our results emphasize the role of hydrodynamic interactions in driving the system towards perfectly in-phase or perfectly out of phase responses depending on the nature of the instability. Specifically, the first bifurcation corresponds to filaments oscillating in-phase with each other. We then extend our analysis to filamentous (line) array and (square) carpets of filaments and investigate the variation of the critical parameters for the onset of oscillations and the frequency of oscillations on the inter-filament spacing. The square carpet also produces a uniform flow at infinity and we determine the ratio of the mean-squared flow at infinity to the energy input by active forces. We conclude by analyzing the bending and buckling instabilities of a straight passive filament attached to a wall and placed in a viscous stagnant flow - a problem related to the growth of biofilms, and also to mechanosensing in passive cilia and microvilli. Taken together, our results provide the foundation for more detailed non-linear analyses of spatiotemporal patterns in active filament systems.

## I. INTRODUCTION

The emergence of rhythmic movements and oscillations in single or arrayed elastic filamentous structures is a common motif in biology. Striking examples are the graceful wave-like beating of distinct frequencies and wavelengths observed in eukaryotic flagella and cilia - organelles found in animal sperm, algae, protozoa, and in respiratory and reproductive tracts [1–5, 7]. Many of the spatio-temporal patterns observed are often planar or near-planar; furthermore, even qualitative aspects of the oscillations are strongly affected by the presence of boundaries and the type of fluid surrounding the cilia [8, 9]. Detailed and innovative experiments have revealed aspects of mechanisms by which different axonemal components are integrated and work together [10–13, 15–21]. Despite the large body of revealing experimental work, the regulatory apparatus that sets the timing and amplitude of the wavelike motions and the physical mechanisms behind the onset of oscillations remain somewhat mysterious and competing theories remain the subject of intense exploration [19, 22–26]. Inspired by these naturally occurring and biologically active filamentous carpets, various synthetic filamentous systems have been proposed that utilize three ingredients - elasticity of the filament, active forces or torques, and dissipation to produce sustained oscillations.

Experimental realizations of these systems typically employ routes that differ both in the means of actuation as well as in their constituents. For instance, active filamentous self-assembling structures are seen to form [27, 28] when ATP is added to cellular extracts comprising of a mixture of microtubules and kinesin motors. Addition of ATP initiates activation of the motors resulting in bundling; the bundled filaments are then observed to oscillate synchronously and sometimes, when arrayed in a spaced line, also form metachronal waves [27, 28]. Similar active filaments are observed in motility assays where elastic filaments (microtubules or actin filaments) interact with molecular motors grafted on a surface (dynein or myosin). These filaments under certain conditions execute planar oscillations (see for instance, [29] and references therein).

A second widely used approach starts from colloidal beads or particles [30–37]. For instance Dreyfus *et al.* [30] have used paramagnetic beads connected together using spring-like biotin-strepdavidin bonds to construct a filamentous elastic structure that can then be activated by external magnetic fields. Sasaki *et al.* [31] report the directed motion of colloidal particles and cater-pillar like motion of self-assembled colloidal chains in a nematic liquid crystal matrix using electrohydrodynamic convection (EHC) rolls. Further adaptions by chemical treating the agents constituting these structures have also been successful in creating synthetic swimmers. Dai *et al.* [33] demonstrate how programmable photosynthetic swimmers can be made by using self-electrophoresis to propel Janus nanotrees. More recently, Nishiguchi *et al.* [34] have demonstrated oscillations in chains of self-propelled asymmetrical colloidal Janus particles fueled by an AC electric field. Interestingly, the elasticity here arises from the field-induced attractive interaction between particles and can be controlled by tuning the frequency of the applied electric field. In these experiments fixing both the position and the orientation of one end of the chain resulted in beating behavior reminiscent of eukaryotic flagella. Further variations incorporating light [35], chemical fields [36], and ultrasound [37] have opened up new avenues in utilizing colloids to create biohybrid and bioinspired swimmers. In a manner mimicking biological cilia and flagella, these synthetic swimmers use the interplay between elasticity, activity (emergent activity due to chemical and light fields or externally actuating fields), and geometry to generate oscillations and movement.

In general, a moving filamentous structure is subject to stresses both along its backbone (tangential stresses) as well as perpendicular to its backbone (normal stresses). Tangentially generated stresses that act along elastic filaments are especially important in the synthetic swimmers just discussed. In such scenarios, the combination of elasticity and stress may induce non-linear buckling instabilities. When a mechanism to dissipate energy is present, such as an ambient viscous liquid in which the filament is placed, oscillations can be automatically generated even in the absence of an external frequency. These buckling induced oscillations have been the subject of recent enquiry. Both continuum and discrete agent based models have been used to investigate the emergence of oscillations in single filaments as well as in moving filament-cargo assemblies [29, 39–46]. Recent analytical and computational studies have focussed on the issue of stability in these follower force driven system. Due to the nature of the follower force, the equations governing the evolution of the filament shape are non-conservative and therefore do not lend themselves to the usual energy minimization approaches to buckling [29, 41, 44, 47, 48]. Therefore, in order to analyze the stability of the filament, one must study the full dynamical response.

Since resistivity theory, which assumes that the hydrodynamic drag per unit length at a point on the filament is simply proportional to the velocity of the element at that point, provides a simple and intuitive way to account for fluid drag on a filament, most studies have employed resistivity theory based approaches in modeling the coupling between hydrodynamic effects and elasticity. For example, recently, Fily *et al.* [29] examined the stability of a straight, slender elastic filament pushing a viscous cargo and subjected a follower force, i.e., an external force acting along its length. Using a one-dimensional elastic filament model, linear stability theory, and fully non-linear computations, they studied the onset of each buckling instability, characterized each buckled state, and mapped out the phase diagram of the system. Fily *et al.* showed that when one of the end of filament is pinned or experiences significant translational but little rotational drag from cargo attached to it, it buckles into a steadily rotating coiled state. When it is clamped or experiences both significant translational and rotational drag from its cargo, it buckles into a periodically beating, overall translating state with the transition to the periodic state occurring via the classical Hopf-Poincare-Andronov bifurcation [49]. Their linear stability analysis was further supported by direct numerical simulations of slightly deformed filaments and consistent with previous Brownian dynamics simulations [39, 40] and analytical results for clamped filaments without viscous cargo [41, 45]. In related work, Ling *et al.* [44] examined the stability of a filament attached to a (virtual) plane wall and acted upon by a follower force. Here, the filament was allowed to also buckle out of plane thereby opening up the possibility of instabilities to a three-dimensional rotating state. These investigators found that the filament first undergoes a bifurcation with a non-planar spinning in a locked curvature. At higher magnitudes of the force, a second bifurcation leads to an in-plane oscillation of the filament. It is noteworthy that both Fily *et al.* [29] and Ling *et al.* [44] used a simple resistivity theory according to which the hydrodynamic force on a filament only depends on the velocity of the filament at that point.

The purpose of the present study is to address the gap in the current theories by using slender body theory to determine more accurate criteria for the stability of single elastic filaments or filament arrays and carpets attached to a sphere or a wall. While resistivity theory gives the correct leading order estimate, of *O*[log(1/*ϵ*)], *ϵ* being the ratio of the filament radius to its length (the aspect ratio), for the viscous drag or traction acting on the filament, in practice its accuracy is limited because of the weak logarithmic dependency of this leading term. To obtain more accurate dynamics of the filaments it is necessary to solve the integral equation for the fluid traction and account for the hydrodynamic interactions. These include interactions between different parts of the same filament, interactions between the filament and a boundary, and inter-filament interactions. Finally, results from the slender body based theory can be used to estimate of the accuracy of the predictions from the simpler resistivity theory based analyses. We should note that the detailed hydrodynamic calculations using slender body theory for a single active filament subjected to a sliding-control based model for active moments in cilia have been carried out recently by Chakrabarti and Saintillan [50]. The present study provides results for another system of active filaments in which the activity is in the form of a follower force.

We first consider in detail the sphere-filament assembly and obtain results for the magnitude of the follower force at which the Hopf bifurcation occurs and the frequency of the sustained oscillations at the onset of bifurcation. It is found that both these quantities vary considerably with the ratio of the sphere radius to the filament length when the sphere-filament assembly is freely suspended. The variations, however, are very small when the sphere is held fixed. Next, we examine the behavior of finite number of filaments attached to a wall and examine how the critical load and the frequency of sustained oscillations depend on the number of filaments and the spacing between the filaments. For multi-filament systems, we find that, although the number of modes of oscillations is equal to the number of filaments, the mode in which all filaments oscillate in-phase with each other occurs at the least load. In other words, all filaments oscillate with the same frequency and in-phase with each other, at least at the first bifurcation from the straight filaments. Following this, we consider a line array of filaments and determine the frequency of oscillations as a function of inter-filament spacing. Finally, we consider a square array of filaments attached to a wall and determine, once again, the frequency and the magnitude of the load at the bifurcation. The array also produces a uniform flow at infinity and we determine the ratio of the mean-squared flow at infinity to the energy input by active forces. Our work generalizes the approach to sphere-filament assemblies (fixed or free to move), as well as to multiple interacting filaments arranged in a linear array or a square carpet - all geometries relevant to applications of the synthetic soft swimmers currently being developed. Besides being of theoretical interest, our analysis will also provide the foundation for more detailed non-linear analysis of possible spatiotemporal patterns in active filament systems. Furthermore, accurate quantification of the parameters required for the onset of instabilities will also aid in the design and control of synthetic swimmers.

The slender body formalism is quite general and can be used to study passive elastic filaments deformed by imposed fluid flows or for a prescribed model of filaments subjected to active moments. Recent studies [51, 52] have addressed flow driven deformation in the context of biofilm deformation. In particular, Guglielmini *et al.* [52] have examined the problem of stability of a straight filament placed in a viscous stagnant flow. Their linear stability analysis showed two bifurcations, the first one corresponding to bending of the filament while the latter one to buckling. These investigators also used the simple resistivity theory for their linear as well as weakly nonlinear analyses. We carry out analysis using the slender body theory allowing for the presence of the wall and thereby provide more accurate estimates for the onset of these two bifurcations.

The organization of the article is as follows. In §II, we consider the sphere-filament assembly. To avoid having to use excessive discretization necessary to resolve the blunt end, we restrict the analysis to filaments that have rounded ends. In §III, we present results for the filaments attached to a wall. Finally, in §IV we consider the case of stability of a filament placed in a quadratic compressional flow near a wall. We conclude in §V by summarizing our results, and suggesting future avenues for exploration.

## II. INSTABILITIES OF AN ACTIVE SPHERE-FILAMENT ASSEMBLY

### A. Slender body theory

The geometry of the sphere-filament assembly is as illustrated by the schematic in Figure 1. The elastic thin filament we consider has length *ℓ* and is comprised of a linearly elastic material with Young’s modulus *E*. The radius of the filament is assumed to be uniform and equal to *ϵℓ* with *ϵ* ≪ 1 except near the end. Furthermore, we assume the filament to be inextensible and unshearable. This allows us to analyze the spatio-temporal deformations using a reduced dimensional form of the Kirchoff-Love equations for the bending of a thin filament [45, 46, 53]. One end of the filament is attached to a rigid sphere of radius *a* = *Aℓ*. Unless specified otherwise in the text, we shall use non-dimensional variables, with distances non-dimensionalized by the length *ℓ* of the filament, forces by the net active force *f*_a_*ℓ*, velocities by *f*_a_/(8*πμ*), and time by 8*πμℓ/f*_a_. Here, *f*_a_ is the magnitude of the active force per unit length acting along the axis of the filament. The assembly is embedded in an incompressible, Newtonian fluid with shear viscosity *μ*.

**FIG. 1.**
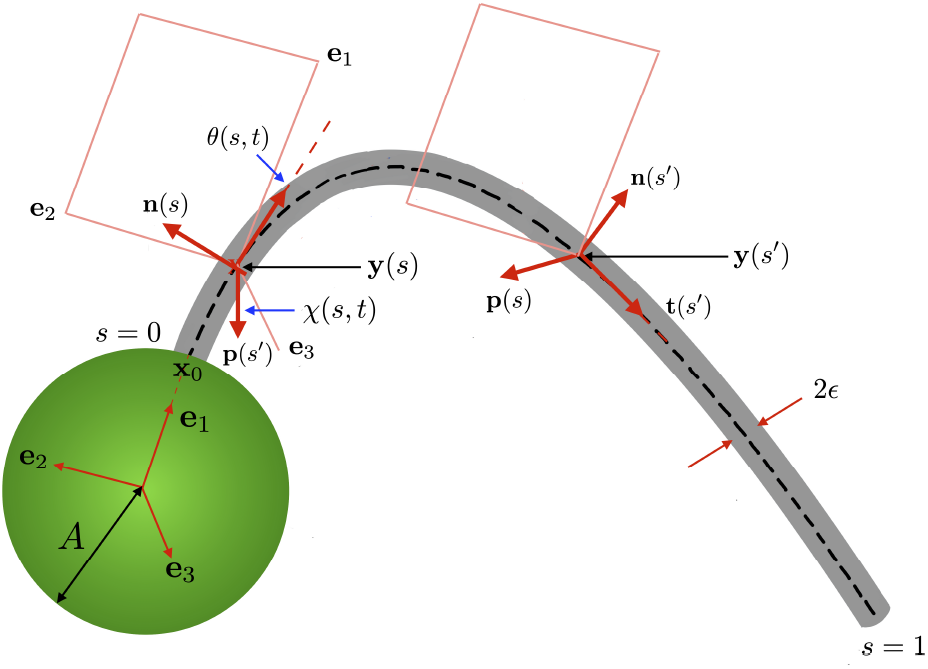
Schematic of the sphere-filament assembly (not to scale) that illustrates the geometry of the system, and orientation of relevant vectors. Here all length scales have been scaled by the filament length *ℓ* (thus the radius of the sphere is *a* ≡ *Aℓ*, and the radius of the filament is *ϵℓ*). The assembly moves in a Newtonian fluid with shear viscosity *μ*. A follower force acts all along the filament - at location *s*, it is directed along −**t**(*s*) and has magnitude *f*_a_ per unit length. We define a (moving) Cartesian reference frame defined by the orthonormal triad (**e**_1_, **e**_2_, **e**_3_) that has its origin at the center of the fixed, rigid sphere. The slender, elastic inextensible filament with a circular cross-section all along its contour length is rigidly attached to the sphere at the point x_0_. Material cross-sections along the filament are parametrized by an arc-length coordinate *s* with the attachment point chosen as the location corresponding to *s* = 0. The other end of the filament *s* = 1 is a free end with force and torque free conditions. Averaging across the cross-section allows us to describe the deformation of the filament as that of a continuous elastic curve. At location *s*, a local material frame is set up by defining the orthonormal triad (**n**, **p**, **t**) located at the centerline position x_*c*_(*s*). Here, **t** is the local tangent vector, and the unit vectors **n** and **p** lie in the plane normal to **t**. This orientation of triad is defined by specifying their directions with respect to the sphere-fixed reference frame using two angles *θ*(*s, t*) (the angle between **t**(*s*) and **e**_1_) and *χ* (the angle between **p**(*s*) and **e**_3_). To simplify the theory, we choose to orient this triad with the principal torsion-flexure axes of the cross-section so that the resulting stiffness tensor is diagonal. As one traverses the filament centerline, the set (**n**, **p**) rotates around the tangent axis. Here, the filament has a circular cross-section, has transversely isotropic properties, is made of a linearly elastic material and has no extensional, shearing or twist deformations.

To specify the shape of the filament relative to the sphere, and the motion of the aggregate, we first define a reference sphere-fixed coordinate system with its origin at the center of the sphere and defined by unit vectors **e**_1_, **e**_2_ and **e**_3_ as shown in Figure 1. The unit vector **e**_1_ is directed along the line joining the center of the sphere with x_0_, the point on the surface where the filament is attached. Next, we choose a base state where the filament is straight and aligned along the **e**_1_ direction. Since filament cross-sections in the undeformed state are circular and the rod is unshearable and inextensible, initially circular cross-sections remain circular. Averaging across the cross-section allows us to treat the filament as an elastic curve (the center-line of the filament) x_*c*_(*s*) with locations (cross-sections) parametrized by the arc-length coordinate *s* measured along the centerline of the filament. The attachment point to the sphere is chosen as the location corresponding to *s* = 0. The other end of the filament *s* = 1 is a free end with force and torque free conditions.

In the following and subsequent discussions, derivatives with respect to the arc-length are denoted by primes for ease of notation. At each point on the centerline, x_*c*_(*s*), we affix a localized coordinate frame defined by the orthonormal triad [**n**(*s, t*), **p**(*s, t*), **t**(*s, t*)] where **t**(*s, t*) is the local tangent vector along the centerline of the filament and the mutually orthogonal unit vectors **n** and **p** lie on the plane normal to the tangent and in the directions of principal axes of inertia of its cross section. (Note that the unit vector **t** should not be confused with scalar *t* which denotes time.) In our case, since the cross-section remains circular, there is freedom in choosing these unit vectors. We specify that for the straight filament **n**(*s, t*) = **e**_2_, **p**(*s, t*) = **e**_3_ and **t**(*s, t*) = **e**_1_. The localized coordinate frame at *s* + *ds* is obtained by an infinitesimal rotation of the coordinate frame at *s*. The deformed state of the axis of the filament is then determined by [45, 53]

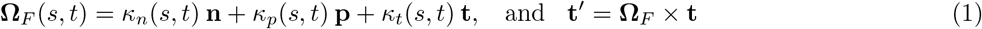

that provides information on the rate at which the triad rotates along the filament. The components of **Ω** are related to the classical Frenet-Serret definitions for the curvature *κ*_SF_(*s*) and torsion *τ*_SF_(*s*) by [54]

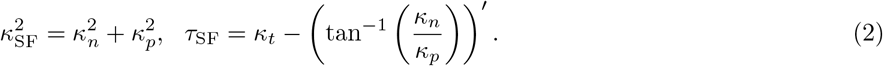

Components of the deformation in the plane perpendicular to t may be extracted using

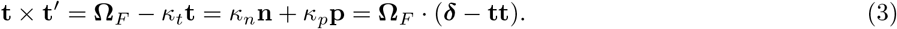

The orientation of the orthonormal vectors (**n**, **p**, **t**) along the filament at every location *s* is defined relative to the sphere-fixed (**e**_1_, **e**_2_, **e**_3_) triad in terms of angles *θ* and *χ* (both functions of *s* and *t*, cf. Fig. 1):

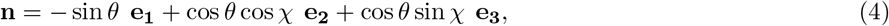

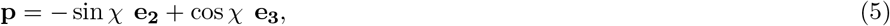

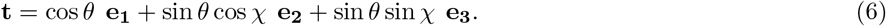

Thus, for the deformed filament, as we traverse the center-line from clamped end to the free end, the set (**n**, **p**) rotates around the tangent axis. The resulting rate of rotation (in space) serves to specify curvatures and torsions. Equations (4)–(6) yield the curvature vector that captures both the bending and direction of the filament at *s* and is related to *κ_n_* and *κ_p_* defined earlier in (1) by

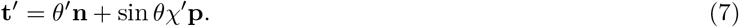

The shape of the filament is completely fixed in the body frame once *θ*(*s, t*) and *χ*(*s, t*) are determined. At each cross section of the filament total force **F** and moment **M** exerted by the hydrodynamic force and the follower force are responsible for bending that deforms the filament. Let *B* ≡ *πEℓ*^4^*ϵ*_4_/4 be the bending modulus of the filament. We then define the dimensionless number

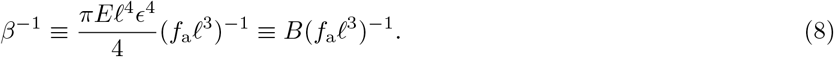

For a filament of given length and stiffness, increasing *β* may be interpreted as increasing the magnitude of the active force. The moment **M** at each cross-section is related to the curvature vector introduced in (1) via Hooke’s law. Ignoring twist, and thus the contributions from component *κ_n_*, the effective internal moment has the form

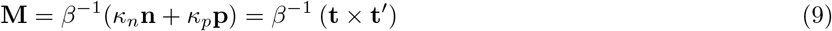

On balancing moments across an infinitesimal element of the filament, we obtain

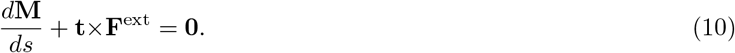

Here, **F**^ext^(*s*) is the total force acting on the filament from a position *s* measured along the center-line of the filament to the end of the filament. Since the force per unit length is scaled by the magnitude of the active force, the active force equals −**t**. Denoting the non-dimensional hydrodynamic traction - the force per unit length exerted by the filament motion on the fluid - by **f** we can then write

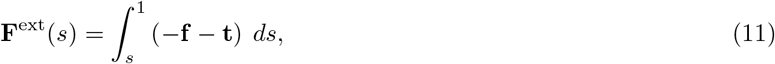

Note that **F**^ext^ is the area-averaged force and is comprised of both an axial force (tension) and a normal force (shear). Combining equations (8)-(11) and using **p** · (**t**×**F**^ext^) = **n**· **F**^ext^ and **n** · (**t**×**F**^ext^) = −**p** · **F**^ext^, we obtain

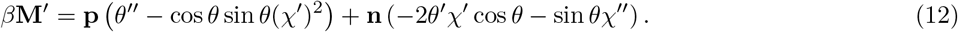

Substituting (11) and (12) into (10) gives

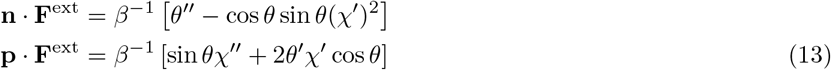

Differentiating above expressions with respect to *s* we obtain the dual set

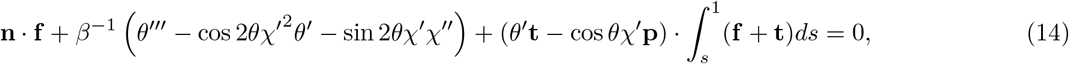

and

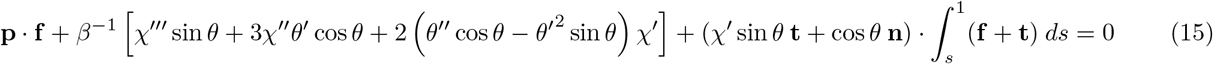

Equations (14) and (15) are Fredholm integral equations of the second kind for the normal components of the hydrodynamic forces and are therefore better suited for numerical analysis than (13), which is an integral equation of first kind for **f**. The unknown function yet to be determined is the hydrodynamic traction **f**. This function and the translational and rotational velocities of the sphere are to be determined by applying the no-slip condition on the surface of the filament, force and torque balances on the sphere-filament assembly, and the moment balance on the filament written in the form of equations (14) and (15).

We shall use the method outlined by Higdon [55, 56] to determine the velocity field and forces on a sphere-filament assembly. The velocity of the fluid satisfies the Stokes equations of motion for a low Reynolds number, incompressible flow. Since the filament is slender (*ϵ* ≪ 1), the velocity at any point **x** *in the fluid outside the assembly or on the surface of the assembly* can be expressed in terms of a line distribution of singularities of Stokes equations of motion along the axis of the filament [55, 56]:

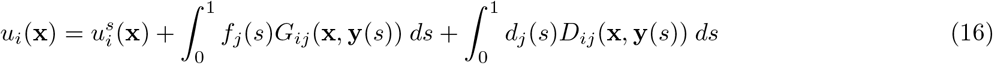

Here, *d_j_* is the source-dipole strength, **y**(*s*) is the position vector of a point on the center-line of the filament at the arc-length *s* (scaled) from the base, and 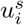 is the velocity induced by the motion of the sphere. *G_ij_*(x, y) is the Green’s function, i.e., the velocity induced at x due to a unit point force applied to the fluid at y in the presence of a sphere. It consists of two parts: 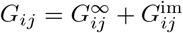 where 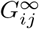 corresponds to the fluid velocity in the absence of the sphere and is given by

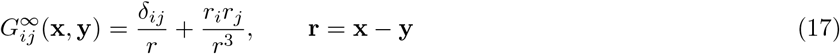

and 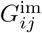 is the velocity due to an image system within the sphere that renders *G_ij_* = 0 for all **x** on the surface of the sphere and at infinity. The reader is referred to Higdon [56] for the expression for 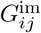. *D_ij_* is the velocity induced by the source-dipole and equals ∇^2^*G_ij_*. The velocity induced by a line distribution of point force normal to the filament axis induces nonuniform velocity on the surface of the filament. Inclusion of the source-dipole term with *d_j_* = *ϵ*^2^(*f_j_* − *t_j_t_k_f_k_*)/2 is necessary to offset this nonuniform distribution. Note that *d_j_* is *O*(*ϵ*^2^) and hence the velocity it induces is significant only near its singularity. The term 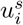 in (16) corresponds to the velocity induced by the translational and rotational motion of the sphere:

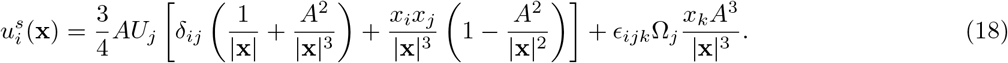

where *U_i_* and Ω_*i*_ are, respectively, the (scaled) translational and rotational velocities of the sphere.

Since the Green’s function we have chosen vanishes at the surface of the sphere, the no-slip condition for the sphere is automatically satisfied. We need to satisfy only the no-slip boundary condition on the filament surface. Since x_0_ is the filament base (cf. Fig. 1), the position vector along the center-line of the filament is given by

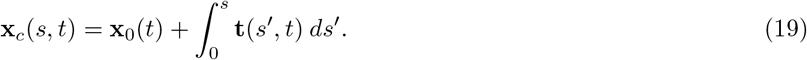

A point on the surface of that element, where the no-slip condition is applied, is given by

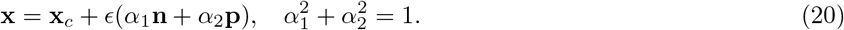

As mentioned earlier, the addition of the source dipole term (*d_j_D_ij_*) in (16) ensures that the velocity induced at the filament surface is nearly independent of the choice of *α*_1_ and *α*_2_. We chose *α*_1_ = 1 and *α*_2_ = 0 in all our numerical analyses.

The velocity of the fluid is evaluated in the body-fitted coordinate system with the unit vectors **e**_*i*_ rotating with an angular velocity Ω_*i*_. The velocity on the surface of the filament is given by

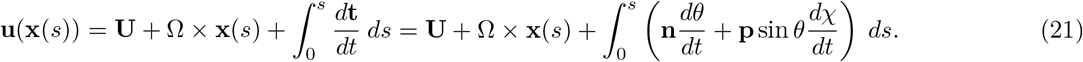

To close the set of equations, we impose the conditions that sum of the active and hydrodynamic forces and torques on the sphere-filament assembly must vanish. This yields the dimensionless constraint equations

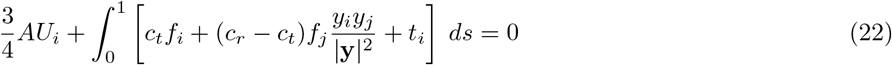

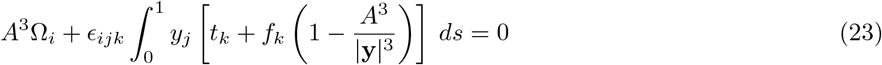

where

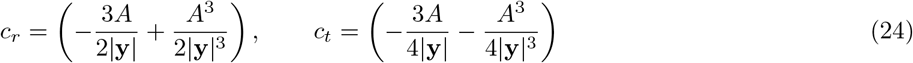

Here, the terms involving *c_r_* and *c_t_* arise from the image forces inside the sphere that are required for satisfying the no-slip boundary condition on the sphere. Likewise, the term *f_k_A*^3^/ |y|^3^ appearing inside the integral in (23) is the torque contribution from the image system inside the sphere [56].

### B. Numerical scheme and boundary conditions

For numerical simulations, we divide the filament into *N* equal elements and approximate *f_j_* by a constant for each element. The force per unit length exerted by the *k*th element on the fluid is denoted by 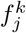 and the center of the *k*th element by 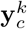. The integral in (16) was evaluated in two parts. The contribution from 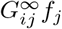 and 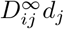 from the *k*th element to a mid-point **x**^*l*^ on the surface of the *l*th element was evaluated using the exact expression given by Higdon [56] while that from 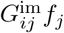 was evaluated using a three-point Guassian quadrature formula. Finally, the contribution from 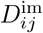 required computing the Laplacian of 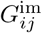. We used a six-point difference formula for evaluating it but found that the results computed with it were essentially the same as those obtained by neglecting the term altogether for *ϵ* = 0.01 or smaller, the cases examined in the present study.

We use trial functions to express the filament shape:

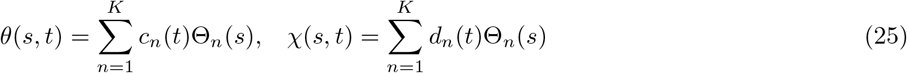

with

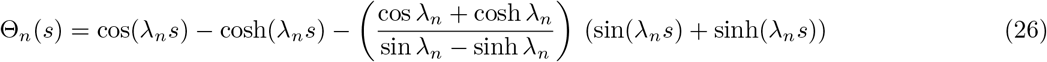

where λ_*n*_ are the roots of the equation cos λ +(cosh λ)^−1^ = 0. These trial functions are chosen such that the boundary conditions *θ*(0) = *χ*(0) = 0 and *θ*′(1) = *θ*″(1) = *χ*′(1) = *χ*″(1) = 0 are automatically satisfied.

Note that setting *θ* and *χ* to be zero at the base of the filament, i.e, at *s* = 0, is equivalent to clamping the filament to the sphere. To observe this we note that close to the attachment point, the lateral displacement of the filament from its straight configuration *W* (in say the **e**_1_ **e**_2_) plane is related to *θ* as *W*′ ~ *θ*. Thus fixing the angle *θ*(0, *t*) = 0 fixes not just the position of the filament but also the slope.

The boundary conditions at *s* = 1 are deduced from (13) by taking **F**^ext^(1) = 0 while those at *s* = 0 are the consequence of choosing the body-fitted coordinates in which the *x*_1_-axis passes through the sphere center and the filament base and not allowing the filament to twist. The functions in (26) are the derivatives of the displacement functions used earlier [44, 45] in the analysis of filaments attached to a wall. Fig. 2 shows the first four trial functions. Application of the no-slip and moments balance conditions on each element and the overall force and torque balances yield a total of 5*N* + 6 equations in 3*N* + 2*K* + 6 unknowns 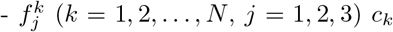, and *d_k_*, (*k* = 1, 2, …, *K*) and *U_j_* and Ω_*j*_. In numerical simulations one can choose *K* < *N* to suppress undesirable oscillations in the filament shape due to higher-order trial functions and solve the resulting unknowns in the least square error sense. For the linear stability analysis carried out in the present study, we took *K* = *N* and solved the appropriate eigenvalue problem.

**FIG. 2.**
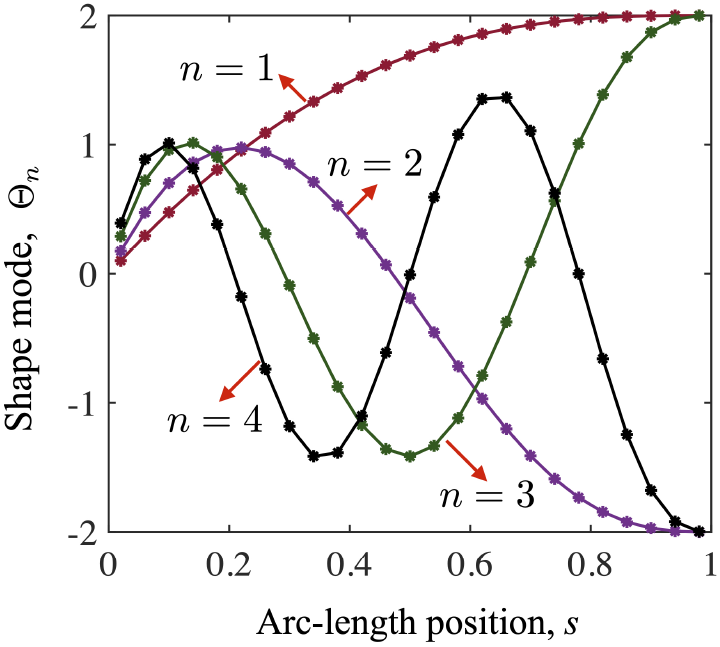
Trial functions Θ*n*(*s*) used for the shape of the filament. The first four functions corresponding to *n* = 1, 2, 3 and 4 are shown.

### C. Base states

We begin our discussion of results by considering first the simple case of steady flows induced by translating and rotating assembly of a *straight* filament attached to the sphere. This is the appropriate base state to consider prior to onset of instabilities. Additionally, this serves to illustrate the convergence of our numerical technique.

Figure 3(a) shows the results for traction as a function of scaled arc-length position *s* when the assembly is acted upon by an active follower force of unit magnitude acting along the axis of the filament. The scaled radius *A* = *a/ℓ* of the sphere equals 0.2, and the slenderness ratio *ϵ* equals 0.01. The wake effect of the sphere reduces the traction on the elements close to the surface of the sphere. Therefore the traction increases with *s*. The numerical results, however, do not converge with increasing *N*, the number of elements used in the calculations. In fact the results obtained with *N* = 40 show considerable fluctuations near the end *s* = 1 even though the translational velocity of the assembly does not change noticeably as *N* is increased from 15 to 40. We therefore further explored modifications that are needed to alleviate the fluctuations and the end effect.

**FIG. 3.**
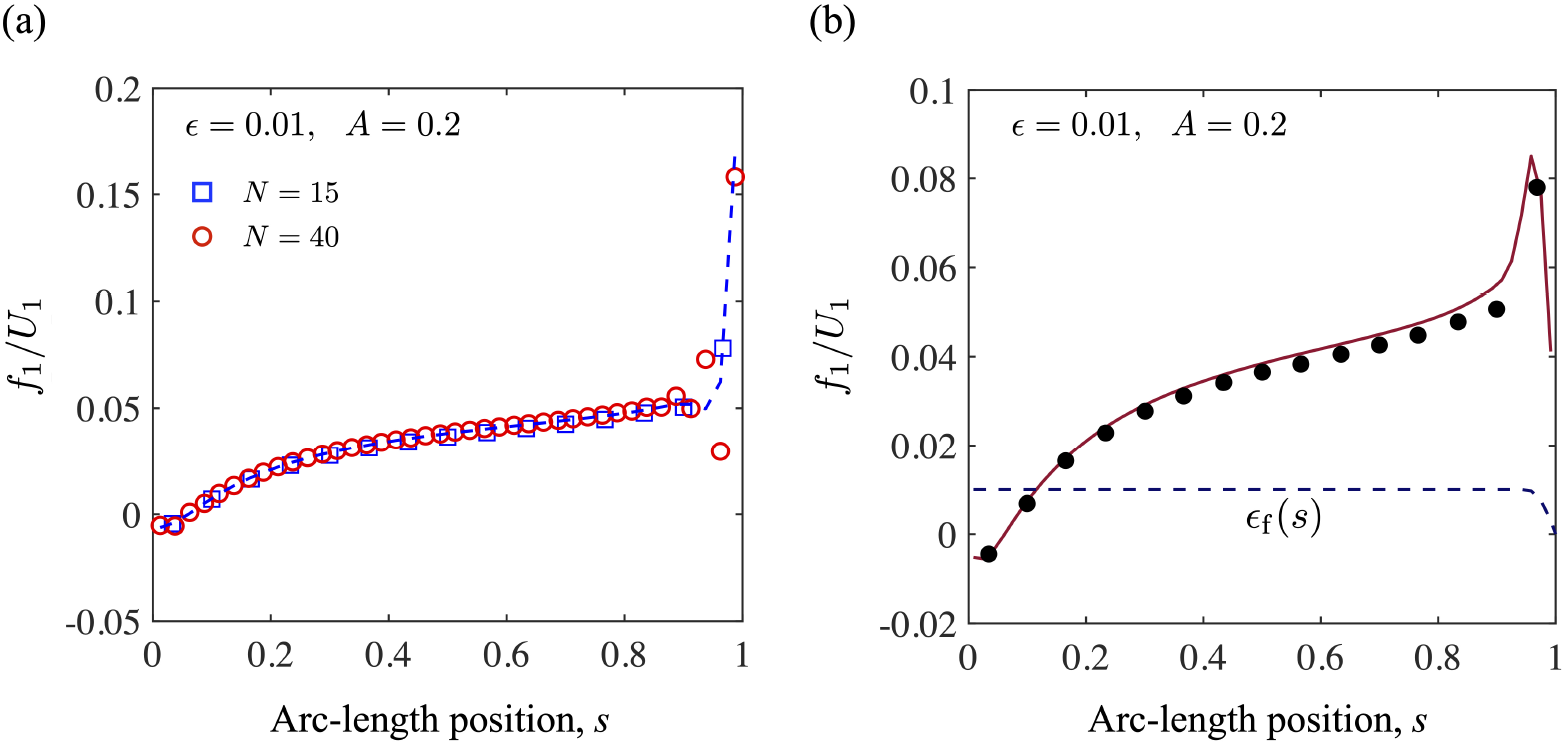
(a) *f*_1_/*U*_1_ for the longitudinal motion of the sphere-filament assembly with = 0.01 and *A* = 0.2. The results obtained with *N* = 15 are indicated by open squares (blue, online); with *N* = 40 by open circles (red, online); and with *N* = 40 and the regularization scheme by a (blue) dashed line. (b) *f*_1_/*U*_1_ for the longitudinal motion of the sphere-filament assembly with *ϵ* = 0.01, *A* = 0.2, and rounded end. The results obtained with *N* = 15 and uniform filament radius are indicated by open circles (black) while those with *N* = 40 and *ϵ*_f_(*s*) as given by (27) are indicated by the solid (red) line.The filament shape used for the latter is indicated by the dashed curve for *ϵ*_f_.

First, we note that (16) is an integral equation of first kind for *f_j_*. It is well known that such equations are generally ill-posed in the sense that they often lead to noisy results because the unknown *f_j_* is entirely inside the integral. It is also known that fluctuations can be suppressed by regularizing the integral equation; in other words by introducing extra terms that penalize gradients in the function that arise due to the ill-posedness. We used a Tikhonov regularization scheme [57, 58] and added a small term *ϵ*(*f_i_* − (5/*N*)^2^*d*^2^*f_i_*/*ds*^2^) to the right-hand side of (16). A central-difference formula was used for estimating the second-order derivatives for all the interior points while the backward or forward difference was used for the elements near the filament ends. The results thus obtained are also shown in Fig. 3(a). We observe that while the regularization scheme suppresses the observed fluctuations in the force density, it still does not produce results that converge for the force density near the end. This second problem arises because our numerical scheme does not account for the no-slip condition on the surface of the blunt end at *s* = 1. In fact, the velocity at (1 + *A*, 0, 0) will be infinite unless *f_i_* approaches zero at *s* = 1. Our numerical scheme only satisfied the boundary condition on the sides of the filament and ignored the no-slip condition on the end plane at *s* = 1 that caps the filament. To resolve this it is necessary to make element length comparable to *ϵ* at least near the end and not apply the force density all the way to the end of the filament. In practice, however, the filament ends are typically not blunt, and therefore added computational effort required to treat blunt end is not worthwhile. The analysis simplifies greatly if one allows instead a rounded end which would ensure that *f_j_* → 0 as *s* → 1. We therefore considered a case of the filament that is rounded near the end *s* = 1 according to

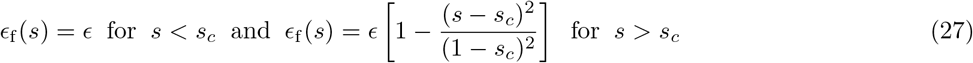

with the cut-off length *s_c_* = 0.95 (cf. Fig. 3b). For such a rounded filament the traction distribution converges and gives *f_i_* → 0 as *s* → 1. The results obtained with the aforementioned regularization scheme and with *N* = 40, which is sufficiently large to obtain reasonably converged results, are shown in Fig. 3b. We see that the traction reaches a maximum near *s* = 0.95 and this maximum roughly equals the results for the straight filament obtained with *N* = 15 that were presented in Fig. 3(a). All results to be presented in the study were obtained with *N* = 15 without rounding the filament or the use of regularization. We expect that the results thus obtained will provide reasonably accurate estimates for the filament that is rounded over approximately last five per cent of its length.

Figure 4a shows the drag coefficient *C*_‖_(*s*) ≡ *f*_1_/*U*_1_ for four different values of *A*; here, the assembly translates parallel to its long axis, i.e., in a direction along the filament axis. The *shadow* effect increases with the increase in *A* and this leads to smaller values of *C*_‖_ for larger spheres. Figure 4b shows the translational velocity of the sphere-filament assembly for a unit applied force for two different values of *ϵ*. The solid lines in that figure correspond to an approximate fit given by

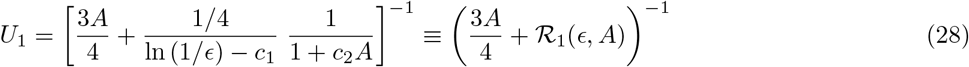

with the constants *c*_1_ and *c*_2_ obtained by a regression analysis to equal, respectively, 0.25 and 5. This formula provides estimates to within 5 per cent accuracy for all the results we computed with *N* = 15 for *A* varying from 0.03 to 5 and for *E* equal to 0.01 and 0.001. Note that 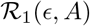 may be regarded as the resistivity of the filament to its motion along the filament length.

**FIG. 4.**
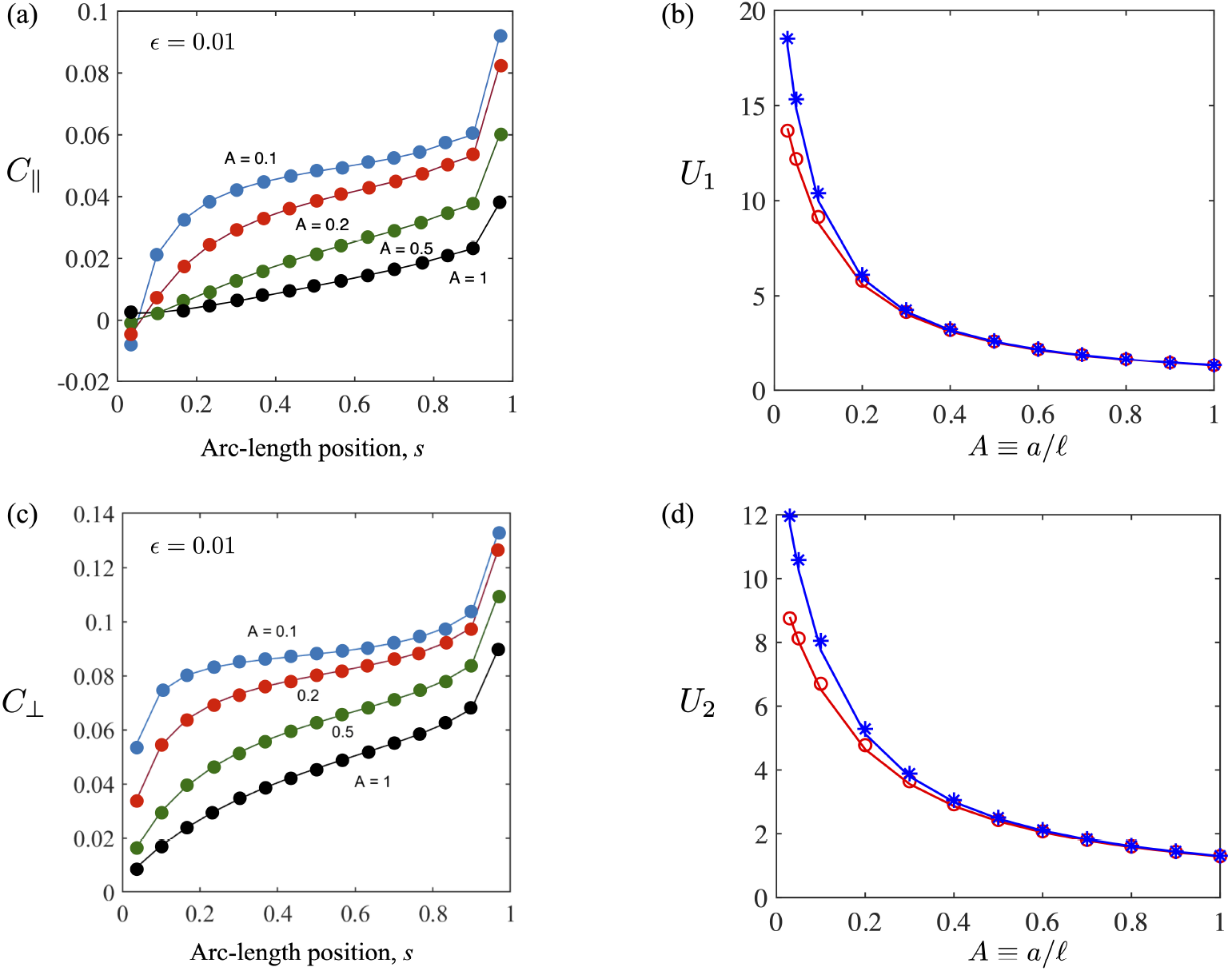
(a) Local drag coefficient, *C*_‖_ *f*_1_/*U*_1_, for the longitudinal motion of the sphere-filament assembly with *ϵ* = 0.01. (b) The translational velocity *U*_1_ of the sphere-filament assembly acted upon by the total force of unit magnitude along the filament axis. The open circles (red, online) and stars (blue, online) are the computed results for, respectively, *ϵ* = 0.01 and 0.001. The lines represent the approximate fit given by (29). (c) Local drag coefficient, *C*_⊥_ ≡ *f*_2_/*U*_2_, for the transverse motion of the sphere-filament assembly with *ϵ* = 0.01. (d) The translational velocity *U*_2_ of the sphere-filament assembly acted upon by a force of unit magnitude perpendicular to the filament axis.The open circles (red, online) and stars (blue, online) are the computed results for, respectively, *ϵ* = 0.01 and 0.001. The lines represent the approximate fit given by (28).

Figure 4c shows the results for the drag coefficient *C*_⊥_(*s*) ≡ *f*_2_/U_2_ for the case when the sphere-filament assembly is translating normal to its axis. As expected, the drag coefficient is seen to be greater than that for the motion along the filament axis. The ratio of *C*_⊥_/*C*_‖_ varies both with *s* and *A*. For example, for *A* = 1, the ratio *C*_⊥_/*C*_‖_ approximately equals 3 at *s* = 0.8 and 7 at *s* = 0.2. Such a large variation in the ratio is not too surprising as the drag along the axis is significantly reduced near the sphere compared with that for the normal motion. The variation in this ratio is smaller for *A* = 0.1 for which it equals about 1.8 at *s* = 0.8, and 2.3 at *s* = 0.2. As seen in Fig. 4d, our results for *U*_2_, for *A* ≥ 0.03 and *ϵ* = 0.01 and 0.001, agree well, within five per cent, with those obtained by the following fit:

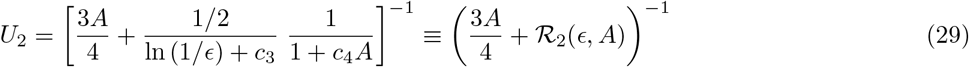

with *c*_3_ = 0.5 and *c*_4_ = 2.5. Equations (28) and (29) show that the net resistance to motion in each case is the sum of the resistances by the sphere and the filament. Once again, 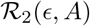 may be regarded as the resistivity of the filament for its transverse motion. Finally, we have also carried out calculations for the case when the sphere-filament assembly is acted upon by a unit torque along the *x*_3_-axis around the center of the sphere. The results for the rotational velocity of the sphere-filament assembly for *ϵ* = 0.01 and 0.001 for *A* ≥ 0.03 agree within five per cent accuracy with the following expression:

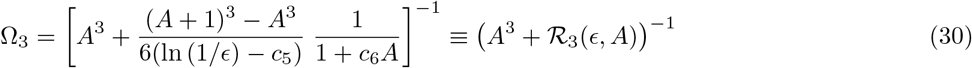

with *c*_5_ = 0.3 and *c*_6_ = 0.45. The effects of shadowing are reflected in the *A* dependence of 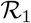, 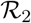 and 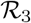.

### D. Linear Stability Analysis

The sphere-filament assembly in the base state that we just examined can become unstable to disturbances that perturb the shape of the filament and deform it. Specifically, the combined effect of the viscous resistance (to both translation and rotation) due to the sphere and the compressive active follower force can make the filament susceptible to buckling instabilities. Here, we explore this possibility and compute parameter values for which this instability is seen.

We begin by examining the effect of small perturbations to the shape of the filament that makes it deviate from its straight shape. We note that the active force is purely axial and along −**t** and deformations initially in the absence of noise are expected to be planar in any plane containing the tangent vector. Let the angles *θ* and *χ* defined in equations (4)-(6) be small, say of *O*(*η*) where *ϵ* ≪ *η* ≪ 1. Terms involving *χ* always appear together with those involving *θ* in the governing equations and therefore do not contribute to the linearized equations up to *O*(*η*). Our linear stability analysis is therefore necessarily limited to small deformations of the filament in the *x*_1_ − *x*_2_ plane. We point our however that equations in §IIA can describe the full non-linear evolution of initially curved shapes which we leave for a future study.

In the small deformation limit, the traction on the filament and velocity of the sphere can be expanded in powers of *η* with the leading-order base state corresponding to the motion of a straight filament acted upon by a force of unit magnitude along the negative *x*_1_-axis. This base state was already examined in detail in section § IIC. We shall denote the traction and the velocity corresponding to this base state by the superscript (0) and the small perturbation quantities without any superscript. The displacement of the filament along the *x*_2_-axis is then denoted in this small deformation limit by

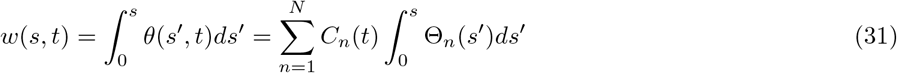

The governing equations for the perturbation quantities are essentially the same as before with minor differences, as described below. The moment equation (14) now reads

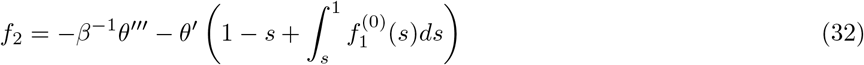

and the no-slip condition for the *x*_2_-component reduces to

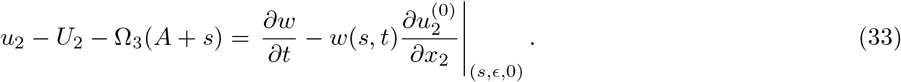

The derivative in the last term on the right-hand side of the above equation is evaluated on the surface of the element at *s*, which corresponds to *x*_2_ = *ϵ* as we have taken *α*_1_ = 1 (cf.(20)) in our base-state computations.

Next, we assume that all *O*(*η*) perturbation variables may be written as exp(*pt*) multiplied by time-independent functions consistent with classical linear stability analysis. For example, we write

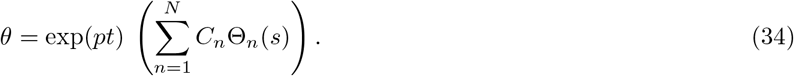

We seek conditions for which the real part of *p* is greater than zero to establish the criterion for the onset of linear instability of the base state.

To determine *p*, we first use the force and torque balance on the assembly to solve for *U_i_* and Ω_*i*_ in terms of *f_i_*. Since we are only considering the two-dimensional motions, we set *f*_3_ = *U*_3_ = Ω_1_ = Ω_2_ = 0 and solve for *U*_2_, *U*_1_, and Ω_3_ in terms of *f*_1_(*s*) and *f*_2_(*s*). Their values are substituted in the no-slip conditions *u*_1_ to determine *f*_1_ in terms of *f*_2_. The moment equation (32) is next used to solve for *f*_2_ in terms of the constants *C_n_* using (25). Substituting next for *f*_1_ and *f*_2_ in the no-slip condition for *u*_2_ in (33) leads to an eigenvalue problem for determining *p* in the form of a generalized eigenvalue matrix equation 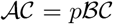 where 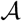 and *B* are *N* × *N* matrices and 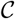 is the vector of trial function coefficients *C_n_*. Solving this eigenvalue equation gives *N* possible values of *p*.

In the problems involving filaments attached to the wall to be considered in the next section, or a sphere that is held fixed, the only modification that is required is that the base state traction and the corresponding velocity are zero. Finally, we note that for the case of a single filament attached to a sphere it is not necessary to solve for *U*_1_ and *f*_1_ as the problem for determining them decouples from that for determining *C_n_* and *f*_2_.

For small *β* all eigenvalues are real and negative indicating that the straight filament is stable to small perturbations when *β* is less than, say, *β*_f_ whose value depends on *A*. For *β* > *β*_f_ (*A*), at least one pair of eigenvalues is complex indicating that the filament will undergo oscillations whose amplitude will decrease with time as the active energy is dissipated away due to viscous drag. Fily *et al.* [29] refer to the critical value *β*_f_ as the flutter point. As *β* is further increased, the real part of the complex pair increases, eventually becoming zero at *β* = *β_s_*(*A*) with further increase leading to positive real parts for the pair. Thus for *β* > *β*_s_, small amplitude perturbations yields oscillations that grow rather than decay with time. This instability where the real component of a complex conjugate eigenvalue pair with non-zero imaginary parts turns positive corresponds to the classic Hopf bifurcation. We shall focus on the onset of these sustained oscillations. At the critical value of *β*, one pair of complex conjugates, for fixed values of *ϵ* and *A*, turns purely imaginary i.e., *p* = ±*iω*. We shall refer to *ω* as the frequency of oscillations.

Figures 5a and 5b show *β_s_* and *ω* as functions of the radius *A* of the sphere and *E*. For *A* > 0.3, the variations in *β_s_* and *ω* are relatively small. The frequencies for *ϵ* = 0.01 are about 30 per cent lower than those for *ϵ* = 0.001 and the critical values of *β_s_* are also lower but only slightly (about 1-2 per cent). However, *β_s_* increases steeply as *A* is decreased and the increase is greater for *ϵ* = 0.01. In fact, for *A* = 0.05 the critical value *β_s_* for *ϵ* = 0.01 (not shown in the figure) is 371, which is more than double that for *ϵ* = 0.001 (*β_s_* = 152). The frequency, on the other hand, always remains smaller for *ϵ* = 0.01 than for *ϵ* = 0.001. The sharp increase in the stability of the straight filaments at smaller values of *A* can be understood in terms of the main terms driving the instability, viz., the quantities inside the parentheses on the right-hand side of (32) which consists of the sum of 1 − *s*, which arises from the active force, and the integral of the base state force 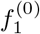. The former is the active compressive force that is the source of bucking instability while the latter, which is induced by the motion of the assembly, acts to suppress the instability. This latter force is proportional to the velocity of the assembly, which in turn is approximately inversely proportional to the radius of the sphere. As a consequence, for smaller *A*, a section of the filament close to the the sphere is under net compression while that near the free end is under tension. For *ϵ* = 0.001, the drag force is smaller and this leads to smaller *β_s_* compared to that for *ϵ* = 0.01 at same *A*.

**FIG. 5.**
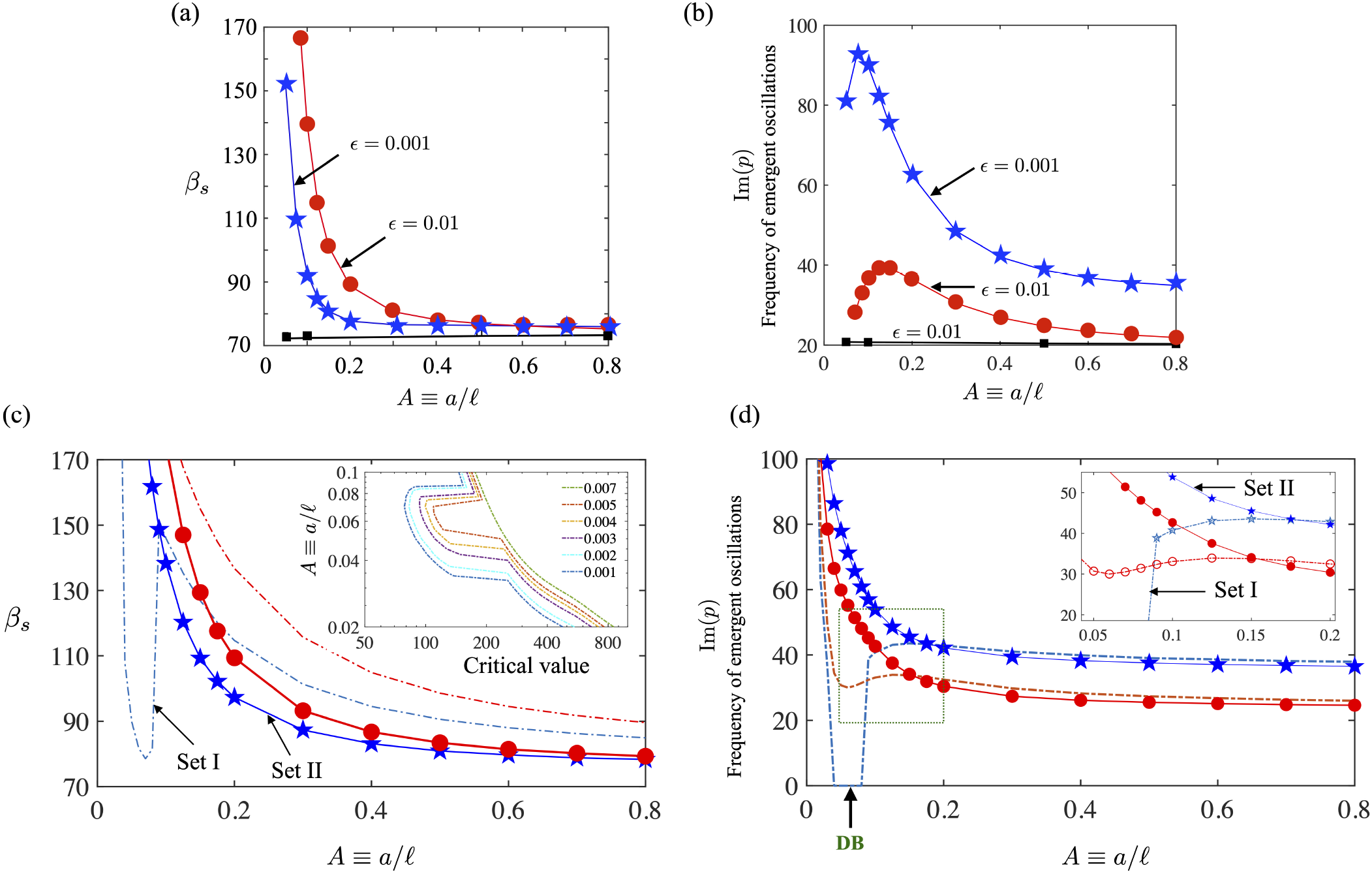
(a,b) Results from the linear stability analysis of the slender body equations for the sphere-filament assembly. (a) Critical values of the non-dimensional active force, *β*, above which sustained oscillations occur as a function of scaled radius *A* ≡ *a*/*l* of the sphere. Filled circles (red, online) and stars (blue, online) correspond to a freely suspended sphere-filament assembly with *ϵ* equal to, respectively, 0.01 and 0.001 while the black squares correspond to the sphere held fixed and *ϵ* = 0.01. Curves/lines are shown to guide the eye. (b) Frequency of sustained oscillations *ω* as a function of *A* at the critical value of *β*. Circles (red, online) and stars (blue, online) correspond to freely suspended sphere with *ϵ* equal to, respectively, 0.01 and *ϵ* = 0.001 while the squares (black, online) correspond to the sphere held fixed and = 0.01. Circles (red, online) and stars (blue, online) correspond to freely suspended sphere-filament assembly with equal to, respectively, 0.01 and 0.001. (c, d) Summary of results from two versions of the resistivity-based minimal formulation described in the Appendix. The first version (Set I, dashed curves) ignores shadowing effects with parameters given by equation (63) in Appendix. The second version (Set II, solid curves and symbols) accounts for them in an approximate way with parameters given by equation (64) in Appendix. Here, red corresponds to *ϵ* = 10^−2^ and blue to *ϵ* = 10^−3^. (c) Critical value of *β* for the onset of instability at which a single real eigenvalue or a complex conjugate pair crosses the real axis. We note that for set II that incorporates the effect of shadowing in an approximate manner, the critical value monotonically decreases with *A* for both values of the aspect ratio. For set I however, this curve for *ϵ* = 10^−3^ does not follow this trend. (Inset) Focusing on set I, we find that the critical value of *β* for instability is non-monotonic in *A* for *ϵ* ∈ (7 × 10^−3^, 10^−3^). (d) Plotting the imaginary component of the critical eigenvalue(s) for both sets I and II shows that the non-monotonic nature arises due to a change in the nature of the bifurcation. For set I, there is a range of *A* for which the critical eigenvalue is a single eigenvalue with zero imaginary component. In this parameter range the bifurcation is a divergence bifurcation (DB) and the emergent solutions are *not* oscillatory solutions. For set II however, instability is always due to a complex conjugate pair and thus the bifurcating branch is a oscillatory solution. (Inset) Close-up of the small *A* region showing the qualitative differences between the two models.

Also shown in Figs. 5a and 5b are the results for the case when the sphere is held fixed and prevented from rotating and translating. In the limit *A* → ∞, this would correspond to the active filament attached to a rigid wall. In this case, the base state force density 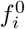 vanishes. Unlike the freely suspended sphere, variations in both *ω* and *β_s_* with *A* are relatively small. *β_s_* varies from 72.3 at *A* = 0.05 to 73.3 for *A* = 0.8 and the the frequency *ω* varies from 20.8 at *A* = 0.05 to 20.3 at *A* = 0.8 for *ϵ* = 0.01. The results for *ϵ* = 0.001 are only slightly different: critical *β_s_* values are, respectively, 74.2 and 74.9 for *A* = 0.05 and 0.8, and the corresponding frequencies are 33.4 and 32.9.

Figs. 5c and 5d summarize results obtained by using two versions of a minimal resistivity theory model (Appendix, equations (57)-(62)) corresponding to two sets of parameters - Set I (equation 63) and Set II (equation 64). Set 1 captures the extra drag and torque due to the sphere on the filament, but does not account for hydrodynamic screening of flows due to the spherical surface. To see if this could be improved upon, we used the results presented in IID that incorporated shadowing effects for a steadily moving straight assembly (equations (28)-(30)) to obtain parameters for a second variant of the resistivity theory (Set 2). In this second model, the effects of the sphere on the filament drag is incorporated indirectly. However, we still do not impose the actual no-slip boundary conditions on the finite sphere. Furthermore, and importantly the effect of the base state fluid flow field on the traction along the moving aggregate is ignored. Thus, in both these resistivity-based models, the compressing force along the filament increases linearly from the free end *s* = 1 to the end *s* = 0 where it is attached to the sphere.

We first discuss the predictions from the minimal resistivity theory without shadowing (set I, dashed curves in figures 5c and 5d). We find that, for *ϵ* = 10^−2^ (red), the critical value of *β* for the onset of instability decreases with *A* for the range shown. The critical eigenvalues are a complex conjugate pair and the instability corresponds to a Hopf bifurcation yielding oscillatory solutions with well-defined frequencies. The critical *β* values compare favorably with the exact slender body estimates (Figure 5a); the frequency dependence on *A* is qualitatively similar. For the smaller slenderness ratio *ϵ* = 10^−3^ (blue), however, we observe qualitative and quantitative differences. Specifically, the critical *β* curve is no longer monotonically decreasing with increasing *A*. Further information about the reason for this difference is obtained by examining the imaginary component of the critical eigenvalue(s). For *A* in the range to 0.088, the base state exhibits two critical points. The first point corresponding to the lower value of *β* arises due to a single *real* eigenvalue crossing the real axis. The bifurcation is thus a simple divergence bifurcation (DB) with small disturbances growing exponentially until non-linear effects limit the amplitude. The second critical point - seen at a higher value of *β* - corresponds to the Hopf-bifurcation. For *A* outside this region, only the Hopf bifurcation exists. Thus the predictions of the minimal resistivity theory are qualitatively and quantitatively different from the slender body predictions.

A possible reason for the lack of the DB solutions in the slender body is due to the boundary condition imposed on the angle *θ* at *s* = 0. To check if this is indeed the case, we next analyzed the predictions from the corrected theory (set II) that incorporates shadowing effects, albeit in a highly simplified form. Our linear stability results are plotted in Figures 5c and 5d (filled symbols and solid curves). We find that incorporating shadowing removes the DB branch - thus for both values of *ϵ*, the straight assembly becomes unstable to oscillatory solutions consistent with the predictions of the slender body theory. The critical values of *β* are also closer to the predictions of the slender body theory. At the same time, the magnitude of the imaginary part of the critical eigenvalues exhibits qualitative differences from the slender body predictions. Specifically, instead of the non-monotonic form of the curves in figure 5b, the predicted curves (solid curves and filled symbols in 5d) are monotonic. To conclude, we find that incorporating shadowing in the manner we did allows us to match slender body predictions in terms of the type of instability and the critical values of *β*. However qualitative differences remain in terms of the emergent frequency of the oscillatory solutions.

Since the derivative of the real part of the pair of complex conjugate eigenvalues at *β* = *β_s_* is nonzero and dissipation provides a mechanism to prevent the uncontrolled growth of the amplitude of the oscillations, we expect stable and sustained oscillations for *β* > *β*_s_. Close to the critical point, 0 < (*β − β_s_*)/*β_s_* ≪ 1 the eigenfunctions corresponding to the critical eigenvalue pair control the shape of the filament deformations. The filament displacement *w*(*s, t*) (i.e., the *x*_2_(*s*) component of the filament center-line) may be determined from the real and imaginary parts of the eigenfunctions corresponding to this complex pair. The result can be expressed (to within a multiplicative constant) according to

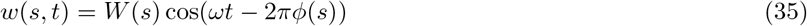

where the phase angle *ϕ*(*s*) is chosen to equal zero at *s* = 0 and *W* (*s*) is the amplitude. Figure 6 shows displacement as a function of *s* over one periodic cycle for the case of a fixed sphere with *A* = 0.8 and for the freely suspended sphere with *A* = 0.05. Figures (7a) and 7(b) show amplitudes and phase angles as functions of *s* for several different cases. The results for freely suspended and fixed spheres are almost identical for *A* = 0.8. Both amplitude and phase angle monotonically increase with *s* for these cases. For freely suspended with *A* equal to 0.3, or smaller, both amplitude and phase angle go through a maximum at an intermediate value of *s*.

**FIG. 6.**
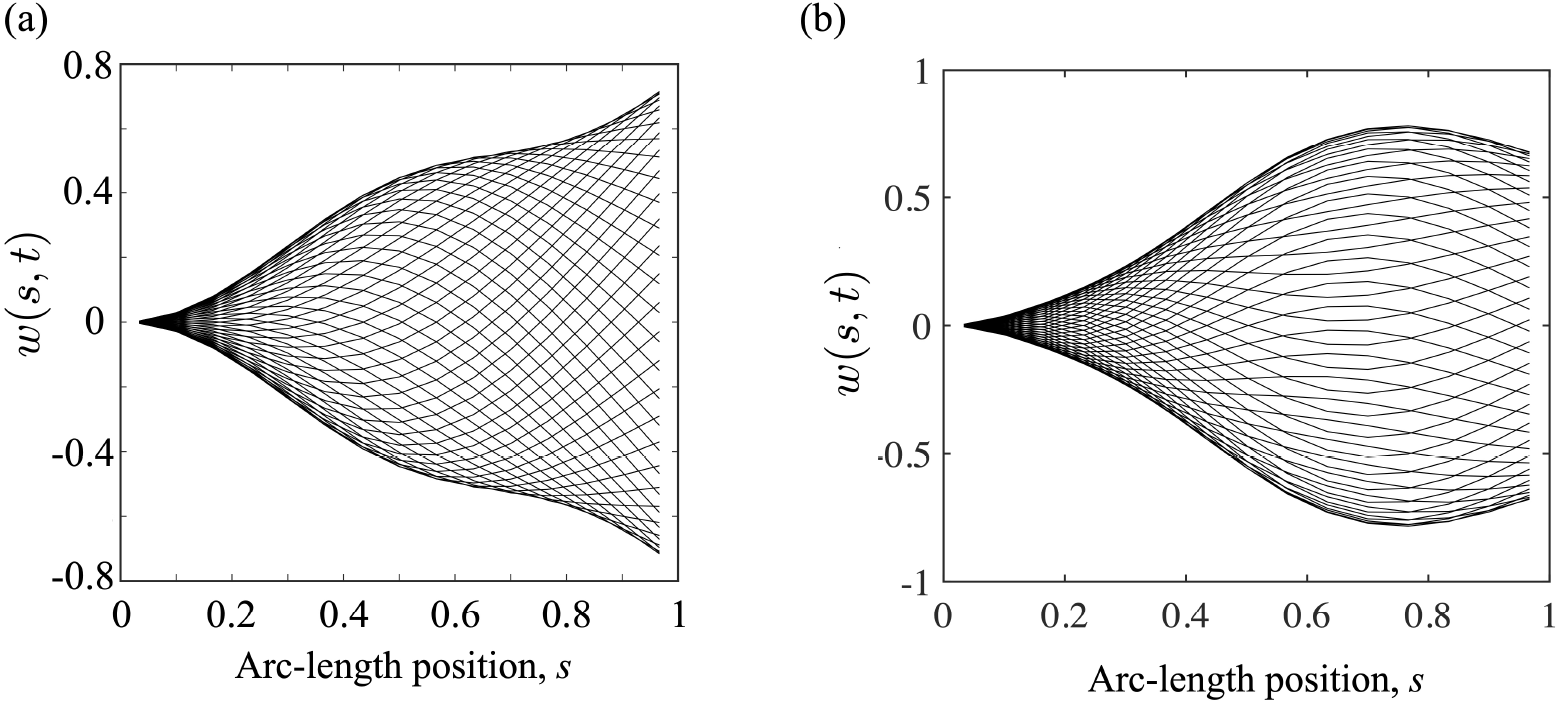
Displacement *w*(*s, t*) - to within a multiplicative constant - of the filament as a function of arc-length position *s* for various times. (a) fixed sphere with *A* = 0.85; (b): freely suspended sphere with *A* = 0.05.

**FIG. 7.**
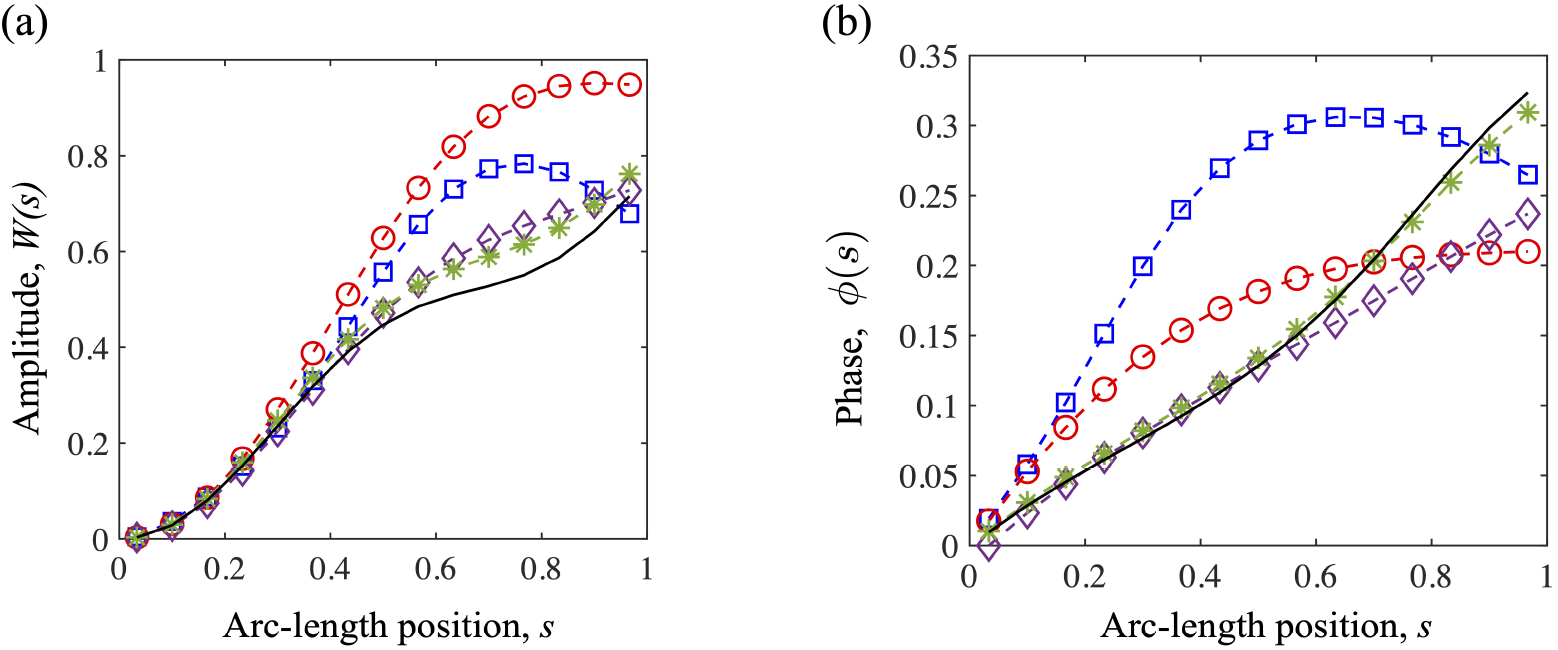
Here we show the (a) amplitude, and (b) the phase angle *ϕ*(*s*), for the following cases: the freely suspended sphere with *A* = 0.05, open squares (blue, online); *A* = 0.1, open circles (red, online); *A* = 0.3, open diamonds (cyan, online); and *A* = 0.8, stars (green, online). The fixed sphere with *A* = 0.8, is shown as the solid black line.

## III. FILAMENTS ATTACHED TO A WALL

We now consider the problem of determining stability of one or more filaments attached to a wall. Each filament is acted upon by a follower force as in the previous section but in addition to hydrodynamic interactions among different parts of the same filament and the wall, we also have to take into account hydrodynamic interactions among different filaments. We shall analyze three cases: (i) a finite number of filaments; (ii) a line array of filaments; and (iii) a square array of filaments.

### A. Green’s functions for the wall-filament geometry

Consider first a single filament attached to a rigid non-moving, no-slip wall positioned on the plane *x*_1_ = 0. The Green’s function must now satisfy the no-slip condition at the wall. Accordingly we write 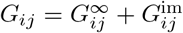 with the latter given by [59, 60]

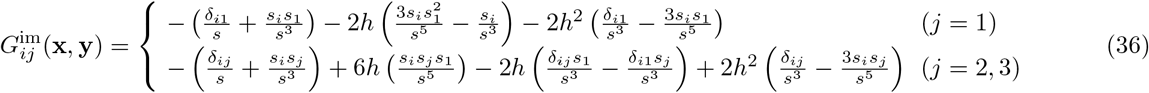

where *h* = *y*_1_ and 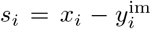 with 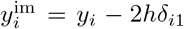. These equations provide a foundation that we will build on to address the hydrodynamically mediated emergent instabilities of multiple interacting filaments. Thus for case (i) of finite number of filaments, the velocity of the fluid is obtained by simply carrying out integration over all the filaments (*α* here denoting the filament index)

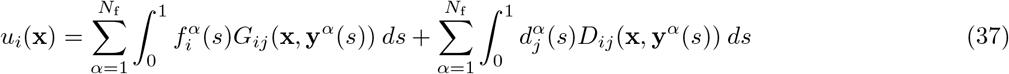

where *N*_f_ is the total number of filaments and **y**^*α*^(*s*) is the position vector of a point along the center-line of the filament *α*. The source dipole strength 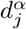 is related to the force coefficient 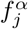 in the same manner as for the case of the sphere.

For case (ii), corresponding to a line array of filaments, the sum over *α* in equation (37) is extended to infinity (−*N*_f_ ≤ *α* ≤ *N*_f_ with *N*_f_ → ∞) and 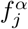 is independent of *α*. It can be shown that *G_ij_*(**x**, **y**) decays as *h*^2^*/r*^3^ for *j* = 1 and as *h/r*^2^ for *j* = 2 or 3, where *r* = |**x** − **y** | and *h* = *y*_1_. Therefore the sum over *N*_f_ in (37) decays as 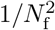 for *j* = 1 and as 1*/N*_f_ for *j* = 2, 3. We computed the sum over *α* for three different values of *N*_f_ and the results were subsequently fitted according to a polynomial 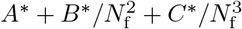 for *j* = 1 and to 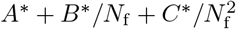 for *j* = 2, 3 to obtain estimate for an infinite array of filaments. It was found that calculations done with *N*_f_ equal to 10, 20 and 30 provided sufficiently accurate estimate for *A*\*, which corresponds to the extrapolated estimate for the array of infinite number of filaments.

The case (iii), corresponding to a square array of filaments that we term a carpet due to the two dimensional coverage of the surface, requires a different expression for the Green’s function. The Green’s function that is spatially periodic in the plane parallel to the wall is derived by Ishii [62] and Sangani and Behl [61]. We write 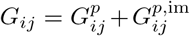 with 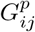 corresponding to the velocity induced by an array of point forces in a plane parallel to *x*_1_ = 0 in an unbounded fluid and 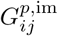 corresponding to the flow induced by its image system in the presence of a wall at *x*_1_ = 0. As shown by Sangani and Behl [61]

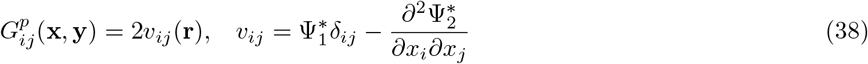

Here, **r** = **x** − **y**, and 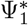 and 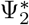 are related to the planar periodic functions Ψ_1_ and Ψ_2_

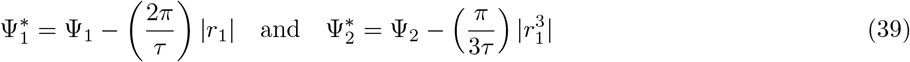

with *τ* being the area of a unit call of the periodic array. The functions Ψ_1_ and Ψ_2_ decay exponentially for large |*r*_1_| so that *v_ij_* increases linearly with *r*_1_ at infinity for *i* = *j* = 2, 3.

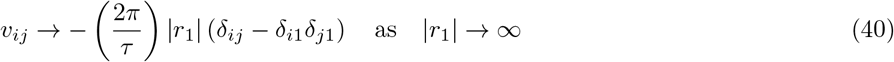

In other words, the velocity induced by point force components parallel to the plane of the array increases linearly at large distances from the plane. It can be shown that the no-slip boundary condition at the wall *x*_1_ = 0 can be satisfied by taking

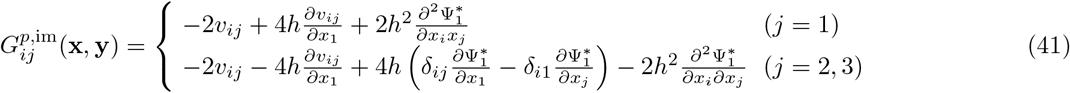

with the functions in the above expression evaluated at *s_i_* = *x_i_ − *y*_i_* − 2*hδ*_*i*1_. Sangani and Behl [61] have described three methods for evaluating the functions Ψ_1_ and Ψ_2_. We used their method I which involes a sum over reciprocal lattice vectors of the periodic array. This sum converges very slowly for small |*r*_1_|. To overcome this problem, we carried out the sum only for reciprocal vectors with magnitude less than 40*/D*, *D* being the size of the array, and approximated the remaining sum by an integral that was evaluated analytically. This analytical integration is possible because, for the case of a straight filament, we only need to evaluate the integral for the special case *r*_2_ = *r*_3_ = 0. For *r*_1_ = 0, Sangani and Behl have recommended using their method II which uses Ewald’s summation technique to obtain fast convergence. We found that for *r*_1_ < 0.05*D* the asymptotic formulas given by these investigators for small **r** provided adequate accuracy and therefore it was unnecessary to use this Ewald’s summation based technique. Note that the linearly increasing part of the velocity induced by an array of point forces of magnitude *f*_2_ is cancelled by their image forces so that the velocity due to an array of point force at a distance *y*_1_ from the wall and its images approaches a constant value equal to 8*πy*_1_*f*_2_/*τ* at infinity. Therefore the flow induced by an array of filaments has the asymptotic behavior

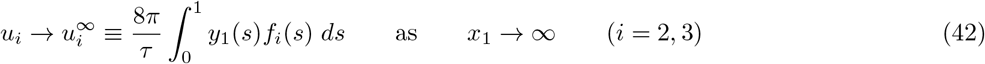

### B. Results: Collective instabilities of a small number of filaments

We first discuss results for the instability of a single filament attached to a wall. The dynamics of a single filament attached to a wall are similar to that attached to a fixed sphere. This is not surprising since the wall is effectively a non-moving sphere with *A* → ∞. The value of the critical load, *β_s_* for Hopf bifurcation varies from 72.18 for *ϵ* = 0.02 to 75.65 for *ϵ* = 0.0001 while the frequency at the onset of instability varies from 16.21 for ϵ = 0.02 to 45.24 for *ϵ* = 0.0001. The following expressions are approximate fits of the numerical results for *ϵ* in this range:

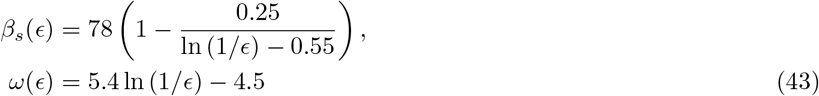

We note that the buckling tendency of the filament due to the imposed follower force is aided by the hydrodynamic drag. Filaments with smaller *ϵ* provide smaller hydrodynamic drag and therefore the required critical load for their instability is greater, i.e., *β_s_* increases as *ϵ* is decreased. The frequency of the oscillations increases as the hydrodynamic drag is decreased.

It is illustrative to compare the values obtained here with previous analyses of the instability of a clamped filament subject to compressional follower forces using closed form analytical equations [29] and discrete Brownian Dynamics simulations [39]. These previous studies differ from the present study in two main aspects. First, the filament is clamped at a point - that is, the wall is virtual and hence cannot affect the hydrodynamic drag on the moving filament. Second, the drag on the filament is calculated using local resistivity theory. Fily *et al.* [29] calculated the critical*β* for instability to be approximately 76.2. A slightly higher value of about 78 was estimated by Chelakkot *et al.* [39] in the weak (but finite) noise regime. Both these values are in good agreement with the values calculated here suggesting that the onset of instability for the single filament problem is controlled primarily by the buckling elastic instability and that the small hydrodynamic drag offered by the filaments affects the critical load only slightly. The effect of hydrodynamic drag is more significant on the frequency of the oscillations at the onset. Since the base state has no flow and no drag (and hence, no influence of the wall), this is to be expected. Note that, the analysis by De Canio *et al.*, [41] is for a point follower load model with the follower force concentrated at the free end rather than being distributed along the filament; the estimated critical value of *β_s_* is therefore different. In terms of frequencies, the simplified analysis of Fily *et. al.* [29] provides the frequency at onset that approximately follows 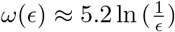 that compares well with the slender body result of equation (43).

We now turn to the more interesting case of dynamics of finite number (*N*_f_ > 1) of filaments attached to a wall. If *N* is the number of trial functions used in the expansion for *θ*(*s*), then the total number of eigenvalues to be determined is *N*_f_ *N*. For sufficiently small *β*, all eigenvalues are real and negative. As *β* is increased there will be *N*_f_ pairs of eigenvalues that will become complex indicating *N*_f_ modes of oscillations. The results discussed below were all obtained for filaments with *ϵ* = 0.01.

We first fix the spacing *D* = 0.3 and study the variation in *β_s_* with *N*_f_. For *N*_f_ = 2, the real part of one pair of complex conjugates becomes zero at *β* approximately equal to 72.5 while the real part of the second pair of eigenvalues becomes zero at *β* = 77. The corresponding imaginary parts, or the frequencies are, respectively, 24.1 and 15.1. The amplitudes and phase angles as functions of *s* for the two filaments are identical at the onset of instability, i.e., at *β* = 72.5, indicating that both filaments oscillate in-phase with each other in this mode. The second mode at *β* = 77 also gives identical amplitudes for both filaments but their phase angles are apart exactly by *π* indicating that this mode corresponds to the two filaments oscillating completely out-of-phase with each other. Since the first mode becomes unstable at the lower value of *β*, we expect the filaments to oscillate in-phase with each other, at least near the onset of the instability of straight filaments. The variation in the critical load and frequency with *D* was also studied and the results are shown in Figures 8(a) and 8(b). These results can be rationalized in terms of the effect of the second filament on the resistivity of a filament. The flow induced by the second filament that is oscillating in-phase with a filament decreases the hydrodynamic resistivity and this leads to lower critical load for the onset of instability and increased frequency of oscillations. This is similar to the effect of decreasing *ϵ* for a single filament, cf. (43).

**FIG. 8.**
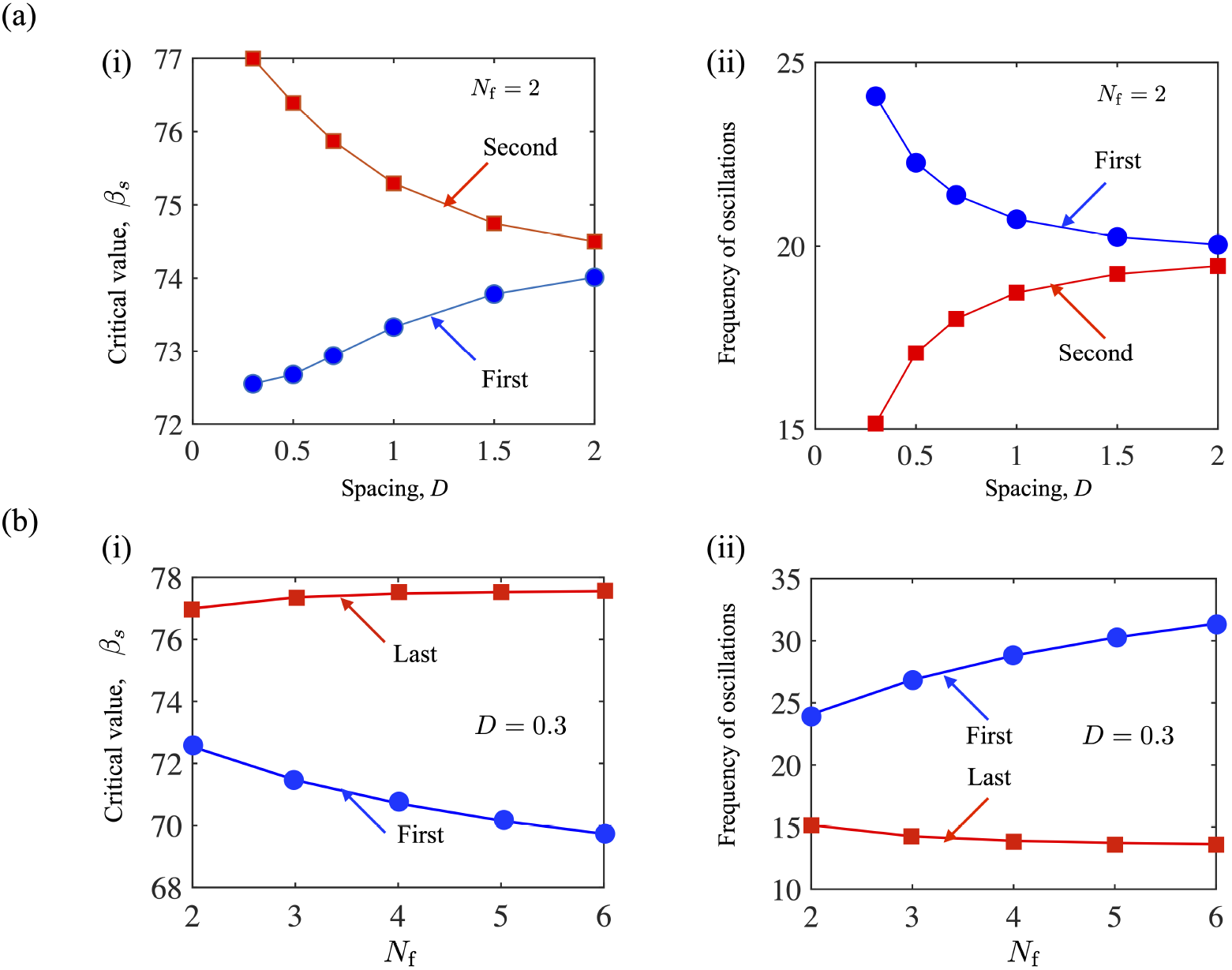
(a) Numerical results for the instability modes for two filaments (*N*_f_ = 2) attached to a wall. (i) The first (circles, blue-online) and second (squares, red-online) critical values of *β* at which oscillatory instabilities emerge are plotted as a function of the spacing *D* between the filaments. The first mode corresponds to the two filaments oscillating in-phase with each other, and the second mode corresponds to the out-of-phase oscillations. (ii) The frequencies of the emergent oscillatory solutions corresponding to these two modes. (b) Numerical results for more multiple filaments. The number of independent modes equals the number of filaments. Critical value of *β*_*s*_ (i) and the corresponding frequency *ω* (ii) at criticality as a function of the number of filaments. There are *N_f_* critical modes and only the results for the first ones that occur as *β* is increased from zero and the last ones are shown. The first mode corresponds to the mode in which all filaments oscillate in phase with each other. The inter-filament spacing *D* is 0.3 and *ϵ* = 0.01.

For *N*_f_ = 3 and for *D* = 0.3, there are three modes of instability corresponding to *β* equal to 71.5 (first), 76.1, and 77.4 (last) with the respective frequencies of 26.9, 17.5, and 14.3. Once again, the lowest *β* mode corresponds to all three filaments roughly in-phase with each other with the amplitude for the middle filament slightly higher than the other two. For the second mode, the amplitude of the middle filament is essentially zero while the two other filaments oscillate completely out-of-phase with each other. Finally, for the third mode, the first and third filaments are in phase with each other with their phase angles differing from the middle one by *π*. The amplitude of oscillations of the middle one is significantly higher than the other two. Once again since the mode corresponding to the smallest *β* for the onset of instability corresponds to all filaments oscillating in-phase with each other, we expect this to be observed in practice.

Even more complex modes are observed for larger values of *N*_f_ but in all cases the first mode to become unstable is the one corresponding to all filaments oscillating in-phase with each other while their amplitudes may differ. The critical values of *β* and the emergent frequencies of the first unstable mode are shown as a function of spacing *D* as well as as a function of *N*_f_ in Figures 8(c). and 8(d) Also shown on the same figures are the critical values of *β* (for oscillatory instability) and the corresponding frequency for the last pair of complex eigenvalues. We note that the first unstable value (corresponding to the lowest value of *β*) decreases with *N*_f_ at fixed spacing (here *D* = 0.3); the associated frequency of oscillations increases with *N*_f_. The trends reverse for the last mode.

We conclude this section by looking at the amplitudes and phase angles of these destabilizing eigenmodes. We choose to focus on *N*_f_ = 4 case. Figure 9 illustrates the results for *N*_f_ = 4 for which there are four pairs of complex eigenvalues. The first mode to become unstable at *β* = 70.7 with *ω* = 28.8 corresponds to, once again, all filaments oscillating in-phase with each other. The amplitudes of the two filaments in the middle are greater than for the other two. Slight differences seen among the middle ones are an artifact arising from boundary conditions being applied at *x*_2_ − *y*_2_ = *ϵ* on each filament, which breaks the symmetry. The second mode to become unstable, at *β* = 75.2 with *ω* = 19.6, corresponds to the first two filaments oscillating completely out-of-phase with the other two while the amplitude of the first and fourth filaments are greater than those of the middle ones. The third mode (not shown), to become unstable at *β* = 76.9 with *ω* = 15.7, corresponds to the first and fourth filaments having the same phase angles but completely out of phase with the two middle filaments. The amplitudes of the outer ones are greater for the middle two filaments. Finally, the fourth mode, to become unstable at *β* = 77.5 with *ω* = 13.9, corresponds to the first and third filaments in-phase with each other and completely out-of-phase with the other two. In this mode, the center filaments oscillate with a greater amplitudes than the filaments at the ends.

**FIG. 9.**
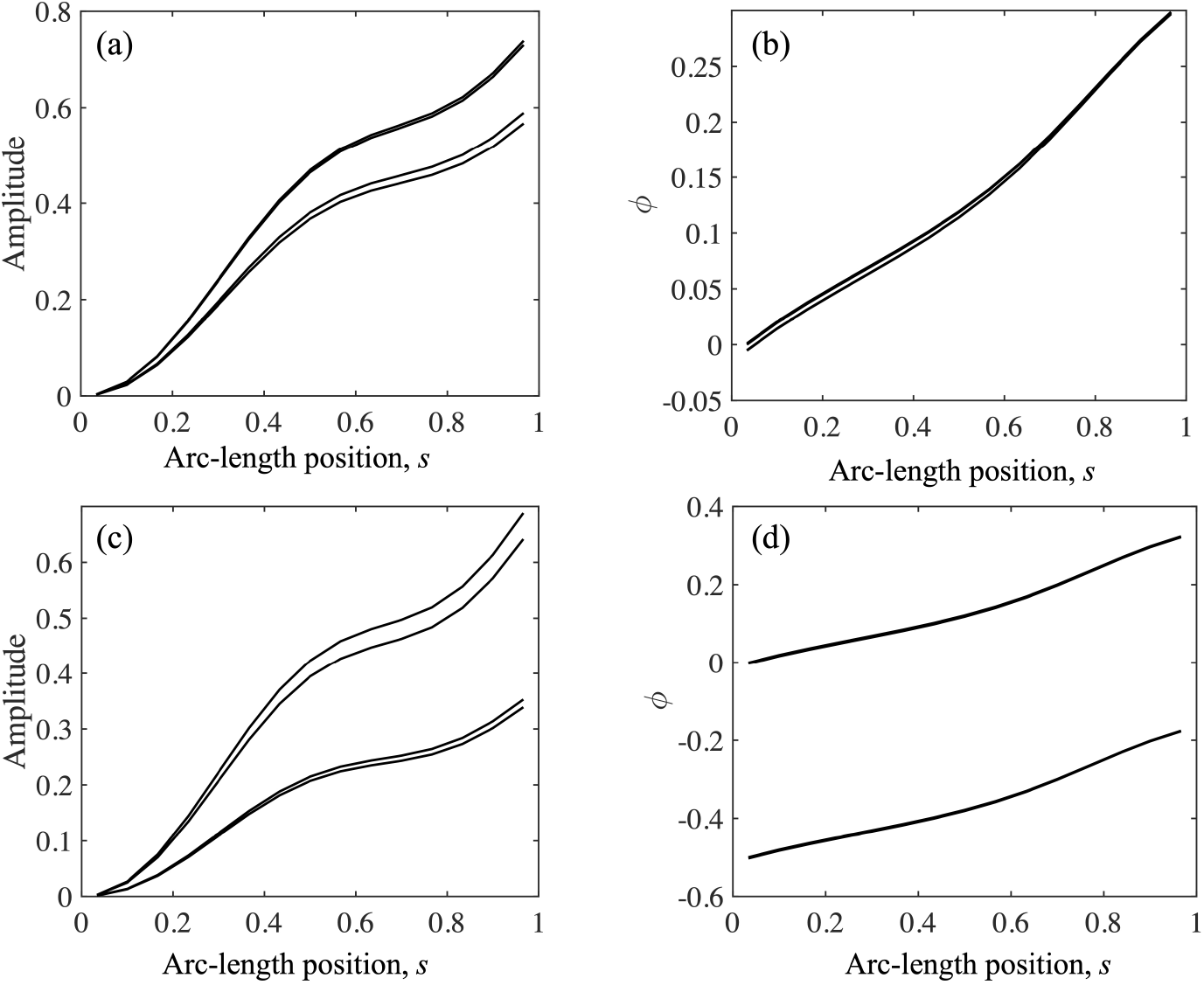
Amplitudes (to within a constant) and phase angles (divided by 2*π*) as functions of arc-length *s* for for four hydro-dynamically interacting filaments. The distance between the filaments is 0.3 and *ϵ* = 0.01. (a) and (b) correspond to the first mode to become unstable while (c) and (d) correspond to the second mode to become unstable. The other two modes that become unstable at even larger values of *β* are not shown.

### C. Instabilities in line arrays and square carpets of active filaments

Next, we present results for periodic arrays of filaments attached to a wall. Figures 10(a) and 10(b) show the critical value of *β* at which the straight filaments become unstable and the corresponding frequency of filament oscillations at the onset of instability as functions of the spacing *D* between the filaments. It is seen that the doubly periodic square array becomes unstable at smaller values of *β* than for the row of filaments having the same spacing and that the frequency of oscillations is higher for the square array than for the line array. Figures 10(c) and 10(d) show the results for the amplitude (to within a multiplicative constant) and the phase angle (cf. (35)) for few selected cases of square and line arrays of filaments. Although the frequency of oscillations and the critical *β* for the onset of instability vary considerably among these cases, we see that their wave pattern is essentially the same, and, in fact, not very different from that for a single filament attached to a fixed sphere with *A* = 0.8. We also note that, in all these cases, the phase angle variation with *s* is nearly linear suggesting that these waves may be easily misinterpreted as traveling waves even though our analysis suggests that the waves at the onset of instability are by nature standing waves that must satisfy the appropriate boundary conditions at *s* = 0 and 1.

**FIG. 10.**
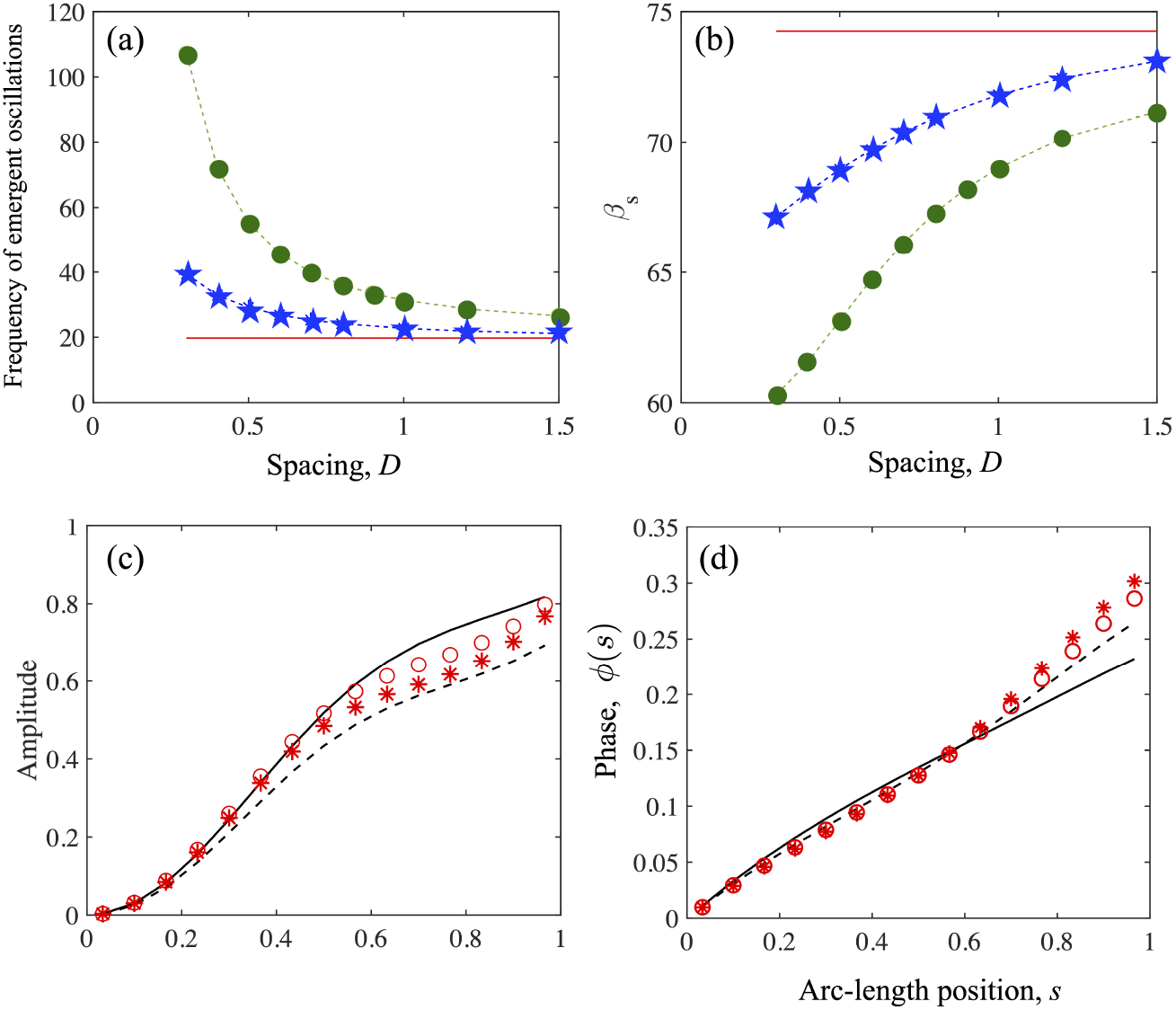
Shown are (a) *β*_s_ and (b) the oscillation frequency *ω* as functions of the inter-filament spacing *D*. Closed circles (green, online) correspond to a square array while stars (blue, online) correspond to a row of filaments in the plane of oscillation. The result for a single filament attached to a wall is indicated by the (red) solid line. (c) Amplitude (to within a constant) and (d) phase angle (divided by 2*π*) as functions of *s* for square and line arrays of filaments at the onset of instability. The solid lines are for a square array with *D* = 0.3; the dashed lines for a square array with *D* = 0.7; the open (red, online) circles and (red, online) stars represent the results for line arrays with spacing equal to, respectively, 0.3 and 0.7.

For the case of square arrays - we term these as carpets in keeping with the literature on ciliary carpets - the oscillating filaments will induce flow at infinity as given by (42). Let us determine the magnitude of this velocity compared to the power input by the active force. To leading order in small amplitudes, the velocity along the *x*_1_-axis of the filament and the force component *f*_1_(*s*) are zero and therefore the power input is entirely determined by the *x*_2_-component of the applied follower force (−**t**) and the velocity of the filament. The *x*_2_-component of the former equals −*θ* while the latter equals *∂w/∂t*. Therefore, the power input by the active force on the filaments per unit area of the square array is given by

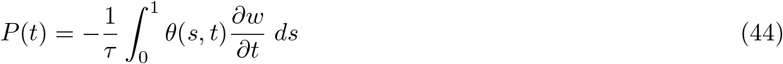

Although the flow at infinity is sinusoidal and hence averages to zero over a cycle, the square of the velocity is not.

We therefore present the results for the ratio

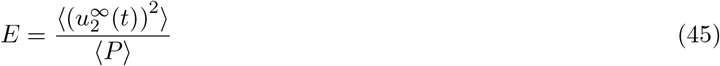

where the angular brackets denote average over one cycle of oscillation. *E* may be regarded as the efficiency of an array of filament in producing the flow at infinity. Note that *P* is non-dimensionalized by 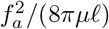 while 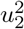 is non-dimensionalized by 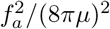 and therefore *E* defined above must be multiplied by *ℓ*/(8*πμ*) to convert it into a dimensional quantity. Figure 11 shows *E* as a function of the inter-filament spacing *D* in square arrays. We observe that *E* varies considerably with *D* with smaller *D* giving higher induced flow. This result is based on power input per unit area. The number of filaments per unit area increases as 1/*D*^2^ which increases more rapidly with the decrease in *D* than the increase in *E* implying that the magnitude of the velocity induced per filament decreases with the the decrease in *D*.

**FIG. 11.**
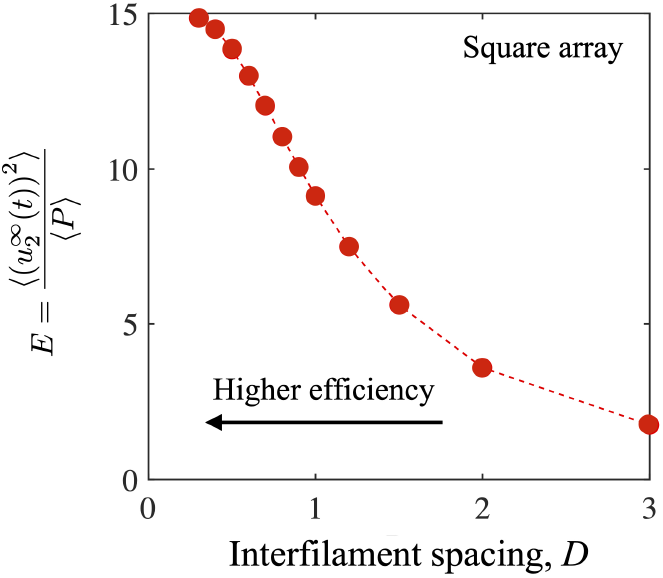
Mean squared flow generated at infinity divided by the mean power input per unit area from the active force acting on the filaments in a periodic array.

## IV. PASSIVE FILAMENT IN A STAGNATION POINT FLOW

In the previous sections, we studied the dynamics and instabilities of elastic filaments subject to follower forces with hydrodynamic interactions arising due to emergent time-dependent behaviour of the system. The slender body formalism is however more general and can be used to study passive elastic filaments deformed by imposed fluid flows. Recent papers [51, 52], and in particular Guglielmini *et al.* [52], have examined in detail the stability of a straight filament placed in a compressive, viscous stagnation point. Their analysis used the leading-order slender body theory without accounting for wall effects or the change in the resistivity with the position of a point on a filament. The formalism presented in earlier sections (§II and §III) is easily extended to treat this problem; we therefore briefly reexamine this problem. Incorporating hydrodynamic interactions with the wall and analyzing the ensuing more accurate slender body theory allows us to obtain more accurate estimates for the onset of the instabilities. The method outlined below may also be used to analyze the elastodydrodynamic deformation of passive arrays and filaments in biological settings such as motion sensing otoliths, stereocilia in ears, and lubricating filamentous aggrecan brushes [63, 64].

We consider a straight elastic filament of length *ℓ* and aspect ratio *ϵ* attached the origin with the wall located at the plane *x*_1_ =0. The filament is subject to a quadratic extensional flow given by

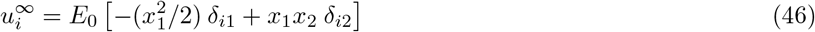

The filament is not acted upon by an active follower force, and so the instability of the straight filament in this case is entirely due to the viscous stresses acting on the elastic filament due to the quadratic extensional flow. After scaling the velocity components by *E*_0_*ℓ*^2^, the critical parameter in the problem can be identified as the parameter *β* = 8*πμE*_0_*ℓ*^5^/*B*, where *B* is the bending stiffness as introduced in §II with 8*πμE*_0_*ℓ*^2^ acting as an effective scale for the force per unit length.

The velocity at a point in the fluid is given by (16) with 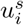 replaced by 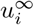. The linear stability of the straight filament is investigated by first solving for the base state corresponding to this flow for a straight filament and then analyzing the equations for small deformations of the filament. The linearized equations and boundary conditions for the perturbation quantities due to slight deformation of the filament can be cast as an eigenvalue problem with the eigenvalues corresponding to the growth rates for the filament deformation.

The numerical scheme we implemented divided the filament into *N* elements and thus matrix to study is an *N* × *N* matrix with *N* eigenvalues. As *β* is increased from zero, we find that one of the eigenvalues first becomes positive at *β* = *β_b_* that depends on *ϵ*. All other eigenvalues remain real and negative. Guglielmini *et al.* [52] referred to this instability as the bending instability. For *ϵ* = 0.01, our computations gave *β_b_* = 122. As *β* is further increased a second eigenvalue becomes positive at *β* = 1396. On increasing the value of *β* to *β* = *β_o_* = 1400, we find that these two eigenvalues become equal (while remaining positive). Further increase in *β* results in these two eigenvalues becoming a pair of complex conjugates with a positive real part. This instability was referred to as the buckling instability by Guglielmini *et al.* [52]. Figure 12 shows the displacement corresponding to these two instability modes at *β* equal to 122 and 1400. Note that for the latter, since the eigenvalues at the critical point equal 0.076, the displacement is proportional to exp(0.076*t*)*W* (*s*).

**FIG. 12.**
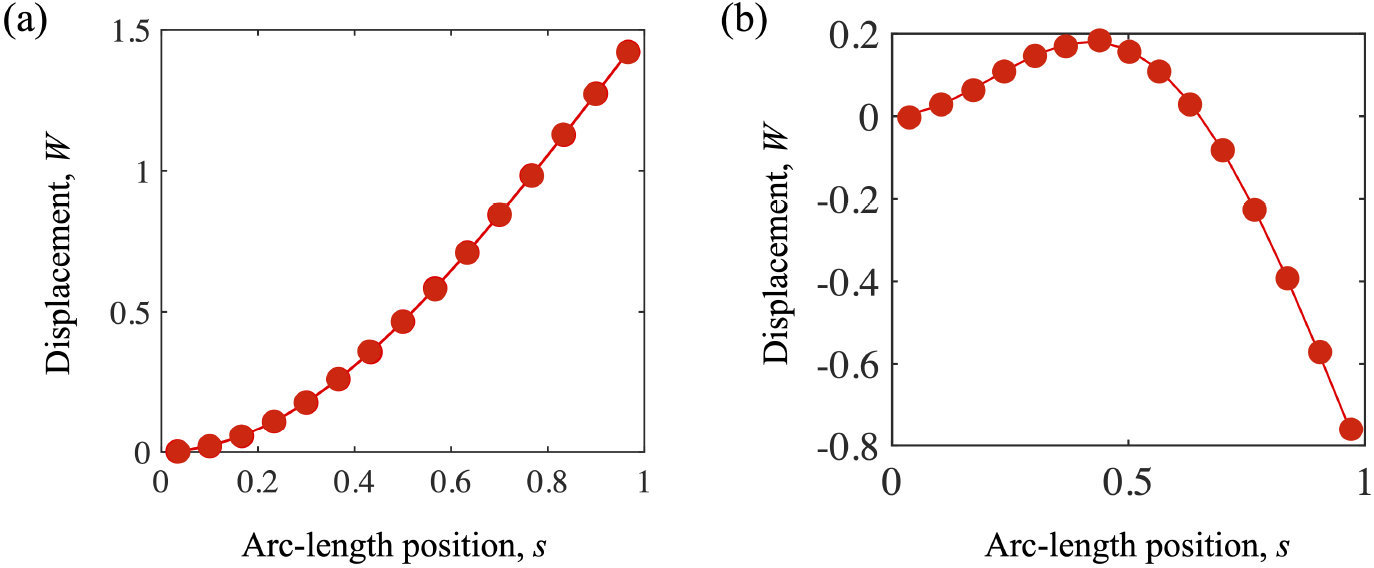
Displacements for the two instabilities for the straight filament with *ϵ* = 0.01 attached to a wall placed in a compressive extensional flow. Displacement as a function of *s* at the onset of bending instability at *β* = 122 is shown in (a) and that at the onset of the buckling instability at *β* = 1400 in (b).

To compare our results with those of Guglielmini *et al.*, we convert their results presented in terms of the parameter they use *η* related to *β* used here by *η* = *β/*(2 ln (1/*ϵ*)). Their analysis gave *η* equal to 9.2 for the bending instability and 250 for the buckling instability. This corresponds to *β* equal to, respectively, 84.7 and 2303 for *ϵ* = 0.01 used in our computations. The former is about 30 per cent lower than our estimate of 122 while the latter is greater by about 64 per cent our estimate of 1400.

It is interesting to enquire if this discrepancy can be reconciled by modifying their leading order estimate of the hydrodynamic resistivity by simply adding an *O*(1) constant. In this case, absent any active follower forces, the key quantity responsible for the instability of the filament to the compressive extensional flow is the tension induced by the base state flow given by

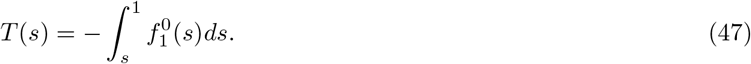

Here, 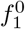 is the *x*_1_-component of the drag force per unit length due to the base flow. In the analysis presented in Guglielmini *et al.* [52], 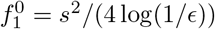, and therefore the tension is given by

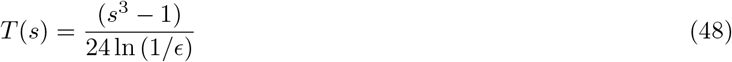

Our numerical results for very small *E*, equal to 0.0001, showed that the computed results for *T* (*s*) are in excellent agreement provided that we modify the above expression by adding an *O*(1) constant so that *T* (*s*) is well approximated by

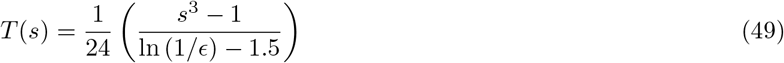

for sufficiently small *ϵ*. Figure 13 shows a comparison of the numerical results for *T* obtained by solving the integral equation for 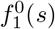 with the above two approximations. We observe that the modified form of the tension *T* (*s*) provides a significantly better match to the computed values even for *ϵ* as large as 0.01 and fully accounts for the effect of the wall.

Figures 14(a) and 14(b) show a comparison of the computed values of *β_b_* and *β_o_* as functions of *E* with the estimates predicted by Guglielmini *et al* [52]. We find that significant discrepancy is observed even if we account for a better estimate of *T* (*s*) by including the above *O*(1) constant. In fact, including the *O*(1) constant only worsens the agreement. The solid lines shown in Fig. 14 represent the fits of the computed results by adjusting both the *O*(1) and *O*(ln (1/*ϵ*)) coefficients:

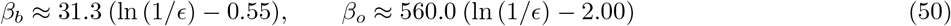

**FIG. 13.**
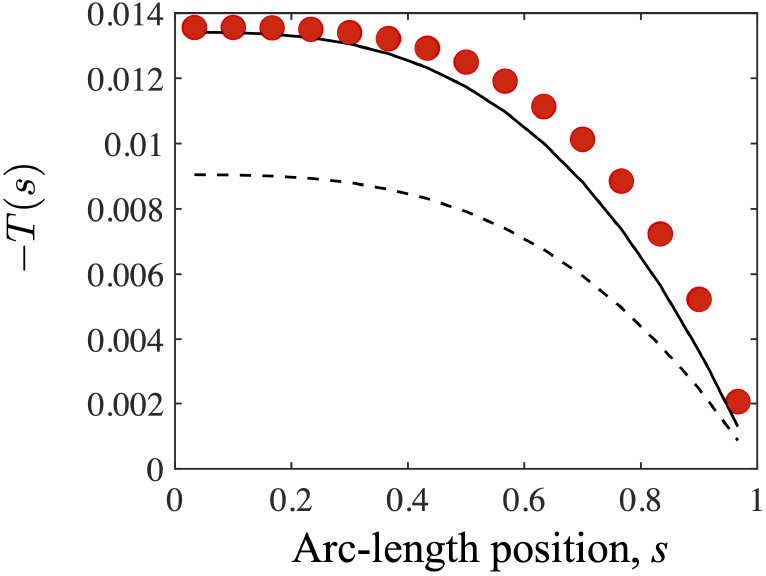
The tension force *T* (*s*) on the filament with *ϵ* = 0.01 attached to a wall in the presence of a compressive extensional flow. The numerical results are indicted by filled circles (red, online) while the approximate values given by equations (48) and (49) are represented by, respectively, the dashed and the solid lines.

**FIG. 14.**
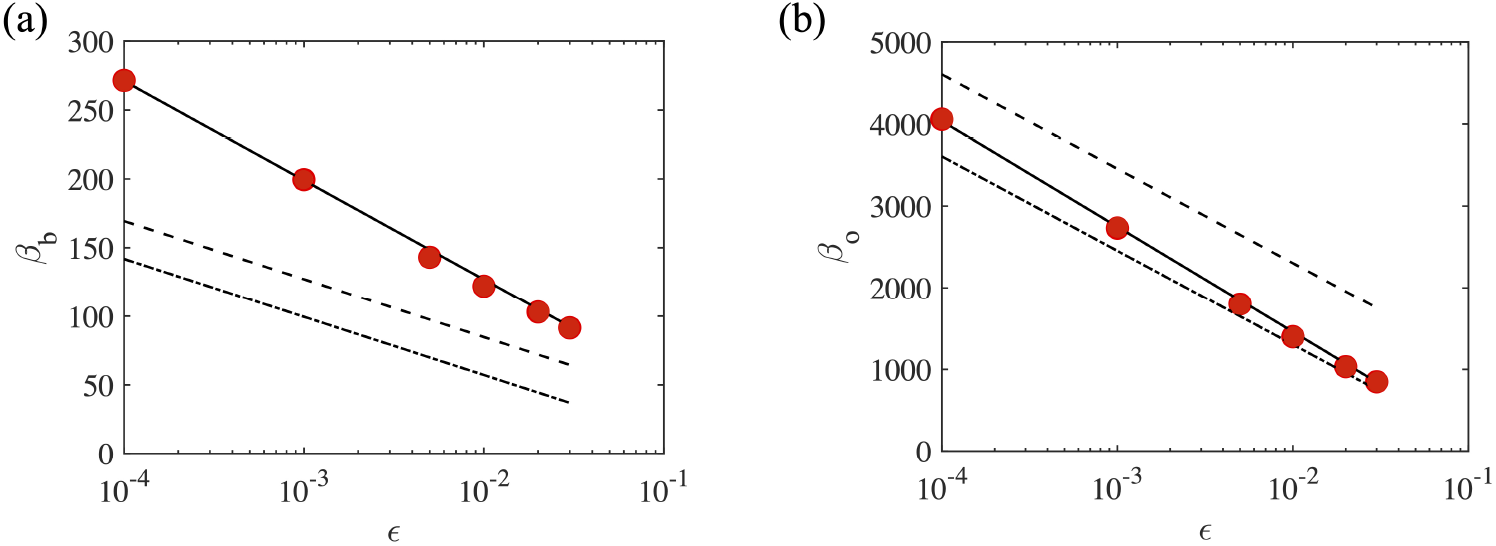
Critical values of *β* as a function of for the onset of (a) bending and (b) buckling instabilities for a filament attached to a wall placed in a compressive extensional flow. The numerical results are indicated by the filled circles while the dashed lines correspond to the result obtained by Guglielmini *et al.* [52] who used leading order approximation for the resistivity. The dotted lines correspond to the prediction obtained by adding *O*(1) constant to their leading log(*ϵ*) terms. The solid lines corresponding to the fit to the numerical results are given by equation (50).

It is not clear why the coefficients of ln (1/*ϵ*) obtained by fitting the numerical results are significantly different from the ones obtained by Guglielmini *et al* [52]. The discrepancies may arise from two sources of approximation. First, it is possible that the form we used for fitting the numerical results, i.e., assuming that the correction to the leading log(1/*ϵ*) term is an *O*(1) constant, may not be adequate for fitting the results as it may be followed by a nonneglible term of, say, *O*(1*/* ln (1/*õ*)) that our fitting procedure does not take into account. We also note that our linearized problem involved the term 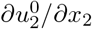 arising from the effect of the base flow on a slightly deformed filament that that has no counterpart in the simplified analysis of Guglielmini *et al* [52]. Therefore, a second probable reason for the discrepancy is that, whereas our analysis accounts for the no-slip boundary condition for the *u*_2_ component by including both the velocity induced by the base flow and the imposed flow (viz. 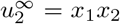), the analysis based on simple resistivity theory by Guglielmini *et al* [52] only accounts for the latter.

## V. SUMMARY AND PERSPECTIVES

In conclusion, we used slender body theory to analyze the linear stability and the hydrodynamic-mediated stable states in active elastic filaments, filament arrays and filament carpets driven by follower forces. The application of slender body theory enables the accurate inclusion of hydrodynamic effects, screening due to boundaries, and interactions between filaments. Our results emphasize that shadowing or screening effects cannot be captured accurately by the local resistivity theory. Specifically, for the case of a freely suspended sphere-filament assembly that mimics synthetic swimmers, the more accurate slender body based analysis provides results that differ qualitatively from the resistivity predictions. The extension to multiple interacting filaments - a small cluster, a linear array or a square carpet - is straightforward in our framework. Our analysis allowed us to investigate the variation of the critical parameters for the onset of oscillations as the frequency of oscillations on elastic, geometric and activity parameters. The square carpet also produces a uniform flow at infinity and we determined the ratio of the mean-squared flow at infinity to the energy input by active forces.

Taken together, our results provide a foundation for more detailed non-linear analysis of possible spatiotemporal patterns in active filament systems. Three possible extensions to further theoretical and numerical analysis are evident. The first concerns the non-linear evolution of the planar oscillatory patterns and the study of interaction between multiple modes of instability post bifurcation, and emergent non-linear, large-amplitude synchronized solutions. In addition to fully stable nonlinear solutions, continuation techniques adapted for time integrators such as the slender body model analyzed in this paper, can identify unstable solutions in both two and three dimensions, as done previously for instabilities in flowing liquid crystal suspensions and in related polymeric processes [65, 66]. Second, our slender body model provides a convenient framework to extend current studies that are focused on applications of Kuramoto theory to interactions between rotating colloids [67, 68] to hydrodynamically interacting filament clusters and arrays. The third extension, which is the focus of our current work, is analyzing out of plane states such as helical waves and rotating states.

## APPENDIX: MINIMAL RESISTIVITY THEORY BASED MODEL

The equations governing the spatiotemporal dynamics of filaments deforming under the action of compressive follower forces and subject to local drag (given by the highly simplified resistivity approximation) have been derived and presented in earlier work [29]. In this work, external molecular motors animated the sphere-filament assembly. There were no singularity distributions to determined since local resistivity expressions were used for the fluid drag. For a moving aggregate, since inertia is absent, the net velocity of translation is obtained by enforcing that the net active force on the aggregate balances the net viscous drag from the fluid. Similarly, the net sum of the active and viscous torque on the assembly is zero (c.f. ESM in [29]).

Here, we specialize the final equations presented there to a sphere-filament assembly restricted to move in the *x* − *y* plane. As introduced in the main text, §II, the assembly comprises of a cylindrical slender elastic filament with length *ℓ*, radius *ϵℓ* (*ϵ* ≪ 1), and bending modulus *B*. The filament is attached to a virtual sphere (point cargo) that exerts a drag equivalent to a sphere of radius *Aℓ*. The assembly moves in a Newtonian fluid of viscosity *μ* due to the action of follower forces of (line) density *f*_a_. Variables are made non-dimensional in the same manner as in §II. Additionally, we denote the (dimensional) viscous drag coefficients (per length) for movement along and normal to the local centerline respectively by *ξ*_‖_ and *ξ*_⊥_.

Treating the filament as an elastic line and restricting deformations to two dimensions, we note that the state and shape of the filament is completely determined by its tension *T* (the area averaged tangential stress), and the angle *θ*. The tension here combines the active and hydrodynamic traction forces and is the magnitude of the effective force introduced in equation (11) in §II. Combining force and torque balances on a differential element of the deforming filament along with geometric constraints, and simplifying the equations for the case at hand (details in [29]), we find the deformed shape to be governed by the coupled non-linear equations

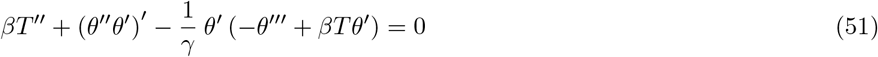

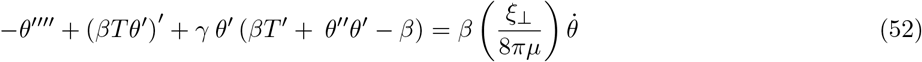

where we have denoted *∂θ*/*∂t* by 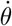 and derivatives with respect to arc-length *s* by primes. Equations (51) and (52) feature three dimensionless parameters *γ* ≡ *ξ*_⊥_*/ξ*_‖_ and *β* ≡ *fℓ*^3^/*B* and the aspext ratio parameter *ϵ* through the exact form of *ξ*_⊥_.

The evolution of the filament’s shape and position is completely specified by solving (51)-(52) subject to boundary conditions. The tail is free, i.e., at *s* = *ℓ* the filament is fully unconstrained, allowing us to write

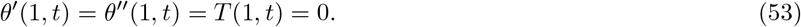

At the head, *s* = 0, we consider an attached sphere that acts as a *point* viscous load. Let the viscous resistance to translation by the head be 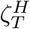 and the viscous resistance of the head to rotation be 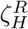. These forces and torques exerted by the (virtual) sphere act on the filament at the attachment point *s* = 0. Defining 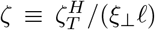 and 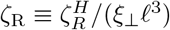, we find the boundary conditions to be,

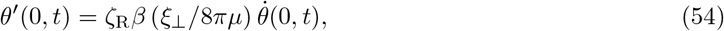

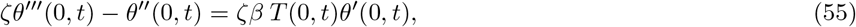

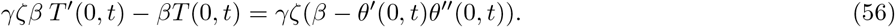

The base state - i.e, the stable configuration under weak loading corresponds to a filament that is straight and subject to follower forces. The sphere-filament assembly therefore translates at constant speed with orientation *θ*_0_ and tension *T*_0_

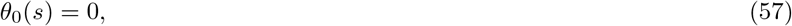

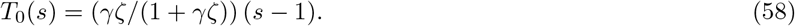

Linear stability of this base state is studied by examining the eigenspectrum of the linear equation

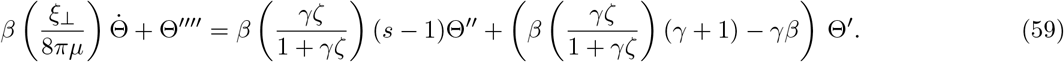

subject to

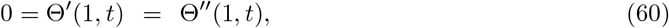

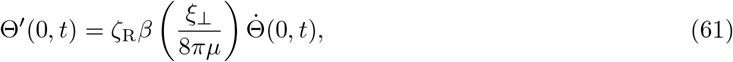

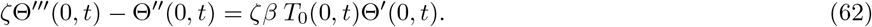

Two sets of parameters were analyzed in this linear stability analysis. In the first - Set I - we ignore the physical presence of the sphere and therefore ignore shadowing effects. The drag coefficients characterizing the normal and tangential resistances to the motion of the filament thus decouples from the sphere size *A*. Furthermore we ignore viscous resistances due to the rotational motion of the filament. These assumptions are found to be quite accurate in modeling the motion of active motor-filament aggregates in motility assays. It is important here to emphasize that in both these minimal models (set I and set II), the base state velocity field is ignored. The fluid velocity field does not enter equations (51)-(64). The base state for the tension - viz *T*_0_(*s*) is thus *a linear function* of position, *s* along the filament, and is not affected by the sphere. For the first set (set I), we set parameters following

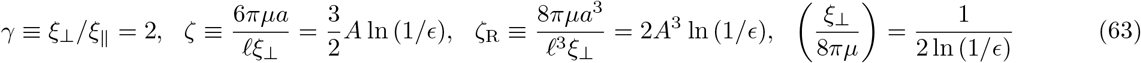

For the second set of parameters - Set II - we utilize results from our slender body analysis that incorporate the first effects of shadowing (equations (28)-(30)). To simplify matters and maintain consistency with the model derived in [29] we here assume that the resistance to the rotational motion of the aggregate about the sphere center can be encapsulated in the rotational resistance of the sphere. Thus for this set we use

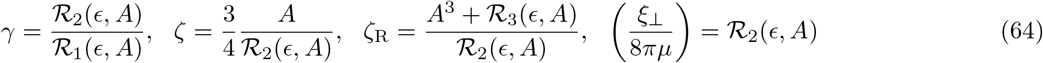

where 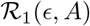, 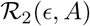 and 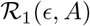 are obtained from equations (28), (29) and (30).

The eigenvalue problem corresponding to equations (56)-(62) was solved using a standard second order scheme finite differences scheme with 51 grid points chosen in the discretization of the spatial coordinate *s*. Care was taken to ensure that the boundary conditions were implemented consistently (see the discussion in ESM [29]). For each set (I or II), substituting 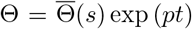 yields the resulting matrix equation whose eigenspectrum was checked. Eigenvalues with positive real parts Re[*p*] greater than a tolerance Tol = 10^−6^ were identified and associated critical value of *β* (equal to *β_s_*(*A, ϵ*)) and their imaginary components Im[*p*] (if this was a complex conjugate pair) were noted. For an oscillatory instability, the frequency of oscillations at onset is then given by *ω* = abs(Im[*p*(*β_s_*)]).

Results are shown in Figures 5(c) and 5(d) for both sets I and II. We note that two modes of instability are observed. The first is a single real eigenvalue crossing the real axis corresponding to a so-called Divergence Bifurcation (DB, in the notation of [29]). The second is when when a real part of a complex conjugate pair goes from being negative to being positive; this bifurcation is termed Hopf-Poincare bifurcation ([49] HB, in the notation of [29]). DB bifurcations in this case eventually lead to the assembly steadily rotating with the filament adopting a shape without inflection points (as seen from fully non-linear solutions [29]). HB bifurcations meanwhile are associated with oscillatory, flutter instabilities with the filament adopting a sequence of oscillatory shapes characterized by a clear amplitude and frequency. The net active force propels the sphere-filament assembly and enables global translation.

